# IRGM/Irgm1 Aggravates Progression of Atherosclerosis by Inducing Macrophage Apoptosis through the MAPK Signaling Pathway

**DOI:** 10.1101/2021.01.06.425662

**Authors:** Shaohong Fang, Song Sun, Hengxuan Cai, Xinran Hao, Xiaoyi Zou, Xin Wan, Jiangtian Tian, Zhaoying Li, Shanjie Wang, Zhongze He, Wei Huang, Chenchen Liang, Zhenming Zhang, Liming Yang, Jinwei Tian, Bo Yu, Bo Sun

**Affiliations:** Department of Cardiology, Second Affiliated Hospital of Harbin Medical University, Harbin 150086, China; The Key Laboratory of Myocardial Ischemia, Chinese Ministry of Education, Harbin 150086, China; Department of Neurobiology, Harbin Medical University; Neurobiology Key Laboratory, Education Department of Heilongjiang Province, Harbin 150086, China; Department of Pathophysiology, Harbin Medical University-Daqing, Daqing 163319, China

## Abstract

**Aims:** Atherosclerosis underlies most cardiovascular diseases, among which acute coronary syndrome (ACS) caused by plaque rupture (PR) often leads to death. Immune-related GTPases (IRGM/Irgm1) have been extensively studied in inflammatory diseases, but their role in atherosclerosis is unclear. Determining how IRGM/Irgm1 promotes atherosclerotic plaque vulnerability will provide information for new biomarkers and/or therapeutic targets.

**Methods and results:** We identified ruptured and unruptured plaques by optical coherence tomography, and found that serum IRGM was highly expressed in patients with ST-segment elevation myocardial infarction in PR. We used ApoE^-/-^Irgm1^+/+^, ApoE^-/-^Irgm1^+/-^ mice and chimeric mice to establish a model of advanced atherosclerosis. The results of pathological experiments showed that Irgm1 caused plaque necrosis. The ratio of neutral lipids and cholesterol crystals increases, while the content of collagen fibers decreases, aggravating the destabilization of atherosclerotic plaques. In vitro, we used multiple approaches to confirm that Irgm1 promotes macrophage apoptosis by promoting the production of reactive oxygen species and activating the MAPK signaling pathway.

**Conclusions:** IRGM may be a potential risk factor for PR. Mechanistic studies have shown that IRGM/Irgm1 contributes to the formation and rupture of fragile plaques. This is partly mediated by the induction of macrophage apoptosis via the MAPK signaling pathway. IRGM may offer new strategies for early treatment of ACS.

**Translation view:** Our findings indicate that IRGM/Irgm1 contributes to formation and rupture of vulnerable plaques. It suggests that IRGM may provide a new target for the early treatment of ACS.

## Introduction

Acute coronary syndrome (ACS), the main complication of atherosclerosis, is the commonest cause of morbidity and mortality worldwide (1). Three main pathological mechanisms lead to ACS: plaque rupture (PR), plaque erosion (PE), and calcified nodules (CN) (2, 3). Historically, research on pathological mechanisms of ACS has mostly relied on autopsy (4). Today, intracoronary optical coherence tomography (OCT) makes it possible to identify the plaque morphologies of patients *in vivo* and probe the pathogenesis of atherosclerotic plaque progression (5). Previously, we used OCT to assess distribution of culprit plaque in ACS patients and confirmed that PR has the highest incidence among the three pathological mechanisms above (6). Vulnerable plaques with thin fibrous caps and large lipid burdens are prone to rupture (7). These findings suggest that increased plaque vulnerability is closely related to PR and the subsequent ACS, and warrants an investigation of the factors and mechanisms that influence vulnerability of plaques. Many clinical trials have suggested that age, smoking, hypertension, chronic kidney disease, and dyslipidemia are associated with plaque vulnerability (8–11). However, molecular mechanisms influencing the vulnerability of atherosclerotic plaques remain unclear.

As the primary immune cells infiltrated in atherosclerotic plaques, macrophages play vital roles in regulating plaque development and stability (18–21). Excessive numbers of apoptotic macrophages are found in vulnerable plaques (22). Macrophage apoptosis causes plaque instability and increases risk of adverse cardiovascular events (23–25). Knowledge of upstream targets that regulate macrophage apoptosis would aid clinical assessment of the risk of vulnerable plaque formation.

The Immunity-related GTPase (IRGM in humans, Irgm1 in mice), belongs to a family of IFN-γ-inducible GTPases (12, 13). IRGM/Irgm1 acts in inflammatory bowel disease, tumors, and other diseases, yet few studies have focused on IRGM/Irgm1 in atherosclerotic disease (14, 15). Previously, we found that Irgm1 influences the progression of atherosclerotic plaques by regulating uptake of oxidized low-density lipoprotein (ox-LDL) by macrophages and the formation of foam cells (16). We have shown that IRGM/Irgm1 promotes the polarization of M1 subtype macrophages and thus aggravates the progression of atherosclerosis (17). These studies indicate that IRGM/Irgm1 expressed in macrophages can promote progression of atherosclerosis and regulate the immune response of macrophages. However, it is not known if IRGM/Irgm1 expressed in macrophages can cause plaque destabilization by affecting apoptosis.

Here, we used OCT to assess morphological features of culprit plaques in patients with ST-segment elevation myocardial infarction (STEMI), and classified patients according to the existence of PR *in vivo*. Serum studies revealed that IRGM was highly expressed in patients with ruptured plaques (the RUP group). Expression of IRGM was correlated with PR in STEMI patients, which can thus be regarded as a potential factor contributing to PR. We clarified *in vivo* that Irgm1 was able to modify phenotype and exacerbate vulnerability of atherosclerotic plaques. Finally, we verified that Irgm1 caused plaque destabilization by promoting macrophage apoptosis. This study provides a new pleiotropic role for IRGM/Irgm1 and unveils novel potential targets for the prevention and treatment of ACS.

## Results

### Serum IRGM is correlated with atherosclerotic PR events in STEMI patients

Initially, we investigated whether serum Irgm1 is associated with atherosclerotic PR in STEMI patients (supplementary Figure 6). Among 85 STEMI patients with analyzable OCT images, PR was the underlying pathology in 52 subjects (61.2%). Figure 1A shows representative OCT images of normal vessels and culprit sites of plaques in RUP and NO-RUP groups. Supplemental Table 1 compares baseline clinical characteristics between RUP and NO-RUP groups. Serum LDL-C and total cholesterol (TC) levels were higher in patients with PR, while no significant differences were observed in other indicators. Plaque characteristics as assessed by OCT are presented in Supplemental Table 2. The prevalence of thin-cap fibroatheroma (TCFA) and lipid-rich plaque was higher, and minimum fibrous cap was thinner in the RUP group.

**Figure 1.**
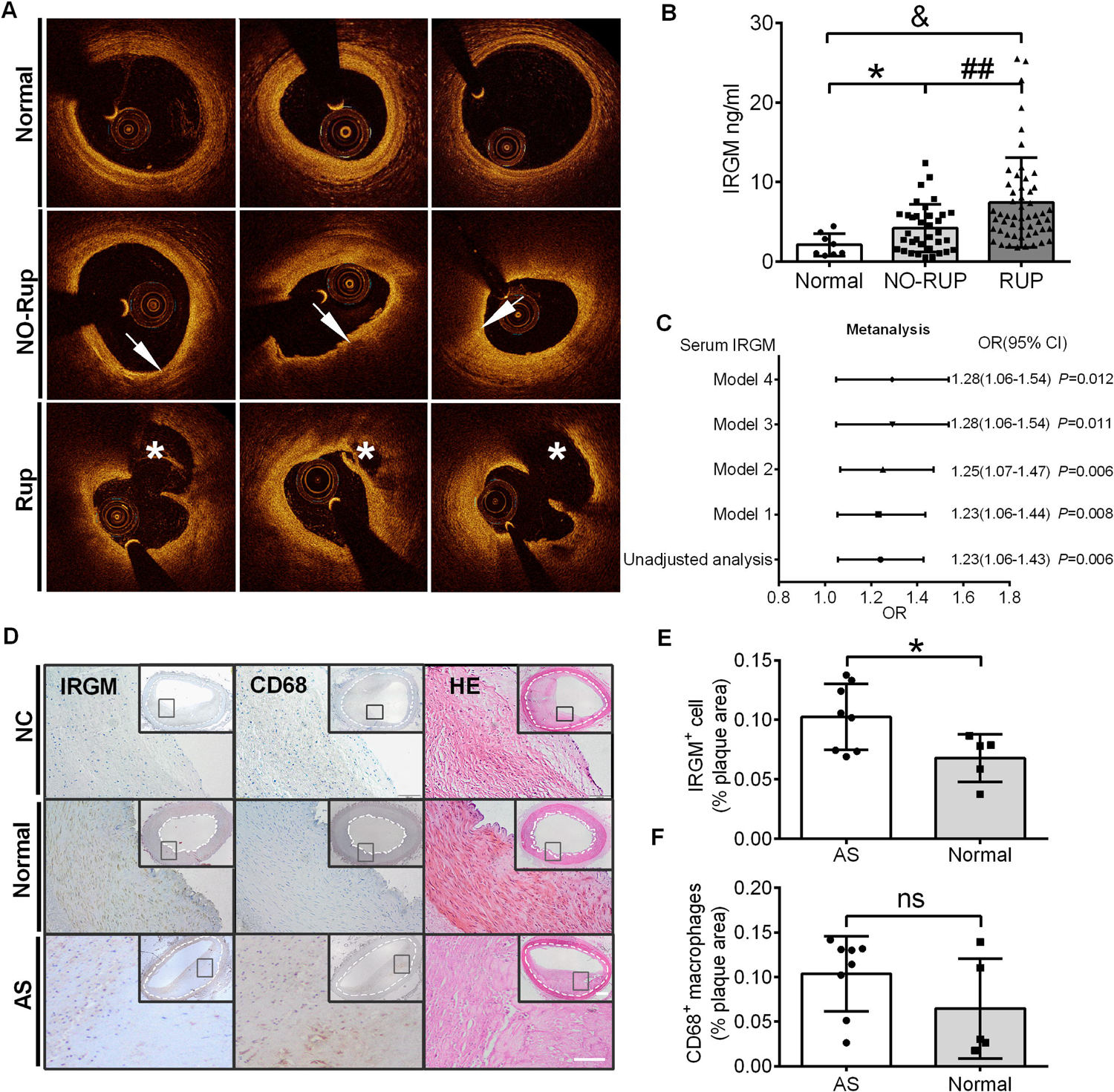
IRGM is associated with atherosclerotic plaque rupture in patients with STEMI, and highly expressed in patients with atherosclerotic plaque. (A) Representative OCT images in normal coronary vessels, NO-RUP plaques and RUP plaques; arrows show TCFA, with FCT < 65 μm, as well as lipid arc > 90°; asterisks represent ruptured cavities at the culprit site. (B) Serum IRGM levels among healthy volunteers, NO-RUP patients and RUP patients. \**p* < 0.05; ##*p* < 0.01; &*p* < 0.05. (C) Adjusted ORs and 95% CIs for plaque rupture associated with serum IRGM in patients with STEMI by 4 models: Model 1, adjusted for age; Model 2, in which Model 1 was further adjusted for hypertension, diabetes mellitus and current smoking; Model 3, in which Model 2 was additionally adjusted for lipid factors (LDL-C and TC) and Model 4, in which Model 3 was further adjusted for peak TnI and hs-CRP. (D) Representative images of the detection of the expression of IRGM and CD68 in atherosclerotic plaques and normal vessels of the human lower limbs. H&E staining is also shown. Scale bars = 100 μm. (E, F) Quantification of the percentage of IRGM^+^μ and CD68^+^ areas in intimal plaques. \**p* < 0.05; ns = no significance.

Serum IRGM was quantified by ELISA, and differed significantly among the three groups. Patients in the RUP group had the highest serum IRGM levels; healthy volunteers had the lowest (Figure 1B).

We next evaluated the ability of serum IRGM level to predict PR in patients with STEMI (Figure 1C). Without adjusting for any factors, IRGM was a strong risk factor, with an odds ratio (OR) of 1.23 (95% confidence interval (CI): 1.06∼1.43, *p =* 0.006). As baseline characteristics differed between these two groups, we adjusted factors including age, hypertension, diabetes mellitus, current smoking, lipid factors, pea troponin I (TnI) and high-sensitivity C-reactive protein (hs-CRP) in four different models. After each increase in the adjustment factors of the models, IRGM remains an effective predictor of PR, with an OR of 1.28 (95% CI: 1.06∼1.54, *p =* 0.012) in model 4. Thus, serum IRGM potentially predicts atherosclerotic PR events in subjects with STEMI, independent of LDL-C and TC levels.

### IRGM is highly expressed in patients with atherosclerotic plaque and positively correlated with CD68

Immunohistochemistry was used to observe the expression of IRGM and CD68 in intravascular intimal plaques. The results showed that expression of IRGM and CD68 in atherosclerotic plaques and their proportion in plaques were significantly increased (Figure 1D–1F). We analyzed the correlation between the expression of IRGM and CD68 in plaques (Supplemental Figure 2F) and found that expression of CD68 and IRGM was positively correlated. Together, these results indicated that IRGM was highly expressed in atherosclerotic plaques and mostly exists in CD68^+^ macrophages within the plaques.

### Irgm1 can affect atherosclerotic lesion phenotype in vivo

To investigate whether Irgm1 can affect atherosclerotic lesion phenotypes *in vivo*, we first distinguished ApoE^-/-^Irgm1^+/-^ and ApoE^-/-^Irgm1^+/+^ mice by genotypic analysis (Figure 2A).

**Figure 2.**
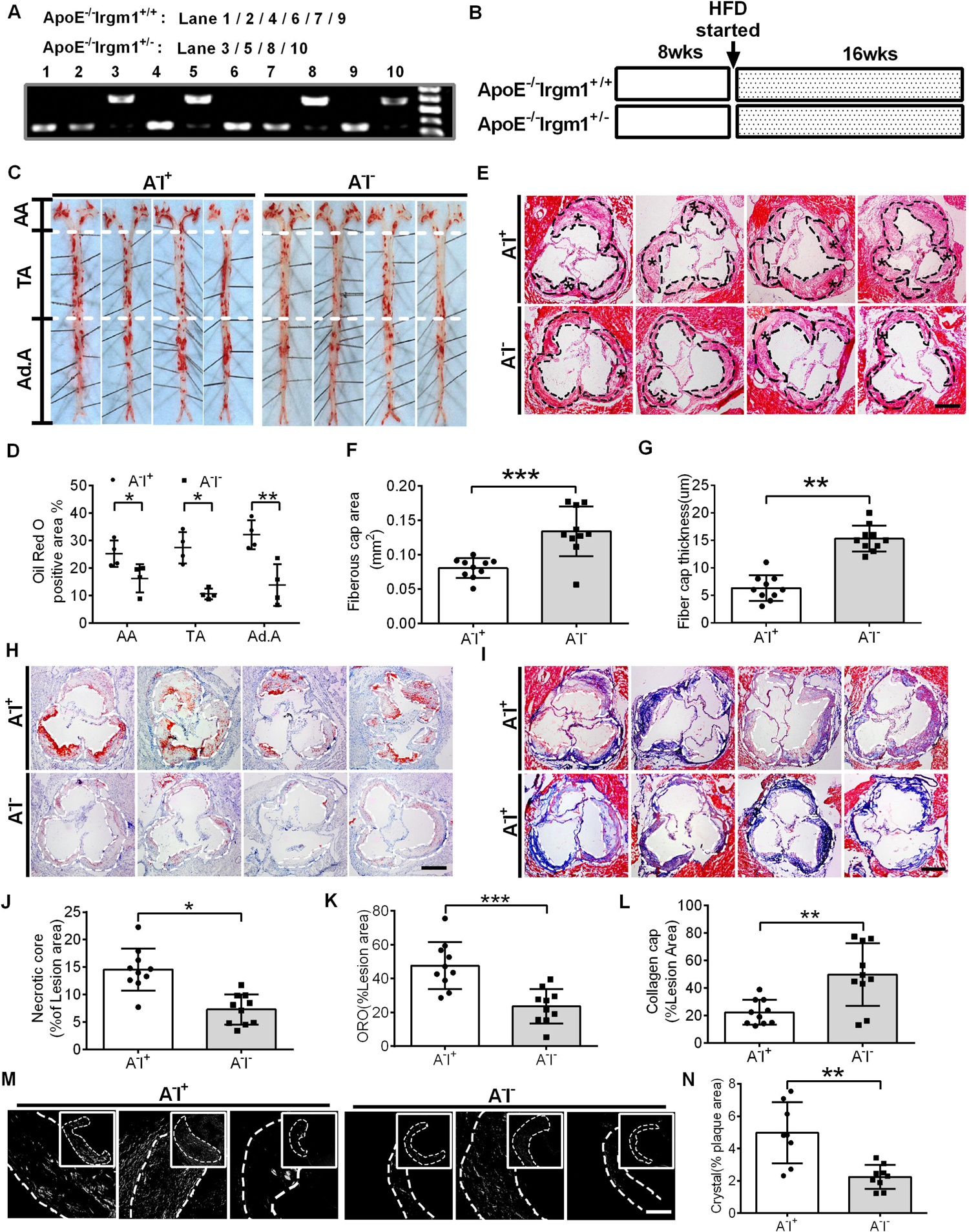
Irgm1 can affect atherosclerotic lesion phenotype in vivo. (A) Genotype analyses for IRGM; Lane 1, 2, 4, 6, 7 and 9 represent ApoE^-/-^Irgm1^+/+^ mice, Lane 3, 5, 8 and 10 represent ApoE^-/-^Irgm1^+/-^ mice. (B) Study design for establishment of advancing atherosclerotic plaques in mice. After being given a high-fat diet for 16 weeks, mice were sacrificed and frozen or paraffin slices of tissues prepared for staining. HFD = high-fat diet; wks = weeks. (C, D) Representative images and quantitative analyses of ORO staining in three parts of the aorta: aortic arch (AA) (n = 4), thoracic aorta (TA) (n = 4), and abdominal aorta (Ad.A) (n = 4). (E) Representative images for H&E staining to assess the FCT, necrotic lipid core and plaque areas in the aortic sinus (n = 10). Necrotic lipid cores are denoted by *; black dashed lines indicate the contour of the plaques; scale bars = 200 μm. (F, G) Quantitative analyses of plaque areas and FCT (n=10). (J) Percentage of necrotic core areas in the plaques (n = 10). (H) Neutral lipid was detected in the aortic sinus by ORO staining (n = 10); scale bars = 200 μm. (I) Representative images for the detection of collagen in the aortic sinus as revealed by Masson staining (n = 10); scale bars = 200 μm. (K, L) Quantitative analyses of the percentage of neutral lipid and collagen content in the plaques, respectively. (M) Representative images of cholesterol crystals observed by confocal microscopy (n = 8∼9); scale bars = 50 and 100 μm. (N) Quantitative analyses of the proportion of cholesterol crystals in the plaque. A^-^I^+^= ApoE^-/-^Irgm1^+/+^ mice; A^-^I^-^= ApoE^-/-^Irgm1^+/-^ mice; \**p* < 0.05, \*\**p* < 0.01, \*\*\**p* < 0.001; ns=no significance. Results are presented as the mean ± SD. Statistical analysis: unpaired Student’s *t-test*.

Compared with ApoE^-/-^Irgm1^+/+^ mice, Irgm1 expression was significantly reduced in eleven major organs or cell types of ApoE^-/-^Irgm1^+/-^ mice (Supplemental Figure 3A). After a Western diet for 16 weeks, mice aorta showed positive oil-red O (ORO) staining, a hallmark of atherosclerotic lesions (Figure 2B-C). There was no difference in the positive area of the aortic arch between two groups, while the positive areas in both the thoracic and abdominal parts were significantly reduced in the ApoE^-/-^Irgm1^+/-^ group (Supplemental Figure 3B). The percentage of ORO positive areas in the aortic lumen area in the aortic arch, thoracic aorta, and abdominal aorta in ApoE^-/-^Irgm1^+/-^ mice were significantly reduced (Figure 2C–D). As shown by H&E staining (Figure 2E), ApoE^-/-^ Irgm1^+/-^ mice had a thicker fibrous cap, reduced necrotic core area and necrotic core ratio, although there was no significant difference in lesion areas between the two groups (Figure 2F–G, 2J; Supplemental Figure 3C). Next, we observed the content of neutral lipids in the aortic sinus of mice by ORO staining. We found that the content and percentage of neutral lipids in ApoE^-/-^Irgm1^+/-^ mice were significantly lower than those in ApoE^-/-^Irgm1^+/+^ mice (Figure 2H, 2K; Supplemental Figure 3E). Masson staining results showed that compared with ApoE^-/-^Irgm1^+/+^ mice, the proportion of collagen content in ApoE^-/-^Irgm1^+/-^ mice was significantly increased (Figure 2I, 2L, Supplemental Figure 3F). Cholesterol crystals in the aortic sinus plaque were further examined by laser reflection microscopy, and ApoE^-/-^Irgm1^+/+^ mice showed significantly fewer cholesterol crystals (Figure 2M–N). These results suggest that Irgm1 aggravates the vulnerable phenotype of atherosclerotic plaque *in vivo*.

### Irgm1 induces apoptosis in macrophages

Previous studies have demonstrated that autophagy affects progression of atherosclerosis (26, 27). We first observed the relationship between Irgm1 and macrophage autophagy *in vitro* (Supplemental Figure 1). After stimulating the RAW264.7 cells with ox-LDL for 3 h, the fluorescence signals of autophagosomes and lysosomes combined successfully (Supplemental Figure 1D–E). Unfortunately, a 48-h treatment with ox-LDL blocked the autophagic flux (Supplemental Figure 1F–G). Excessive apoptosis of macrophages is a feature of vulnerable plaques (22). We investigated whether Irgm1 promotes macrophage apoptosis during the progression of atherosclerosis. Compared with ApoE^-/-^Irgm1^+/+^ mice, the proportion of TUNEL-positive cells in ApoE^-/-^Irgm1^+/-^ mice was significantly decreased (Figure 3A–B). Expression of cleaved-caspase3 and cleaved-caspase9 in the plaque of ApoE^-/-^Irgm1^+/-^ mice was also significantly decreased (Figure 3C–E). These results indicate that Irgm1 can induce apoptosis in plaques. Next we assessed whether Irgm1 induces macrophage apoptosis *in vitro*. The qPCR showed that si-Irgm1 knocked down Irgm1 expression effectively (Supplemental Figure 1A). To strengthen our conclusion, we analyzed the effect of si-Irgm1 on the activities of cleaved-caspase3/9. Compared with the si-control group, cleavage caspase3 and cleavage caspase9 activities of the si-Irgm1 group were reduced (Figure 3F-G), consistent with western blotting results (Figure 3H-I). Furthermore, compared with ApoE^-/-^Irgm1^+/+^ mice, apoptosis of peritoneal macrophages in ApoE^-/-^Irgm1^+/-^ mice was reduced (Figure 3G, 3M). Finally, in vivo, we investigated effects of Irgm1 on expression of cleaved-caspase3 and cleaved-caspase9 in macrophages (CD11b). The results showed the co-localization ratio in the ApoE^-/-^Irgm1^+/-^ mice plaques was significantly reduced (Figure 3K–L, Figure 3N–O). In summary, these results confirmed that Irgm1 promote macrophage apoptosis both *in vivo* and *in vitro*.

**Figure 3.**
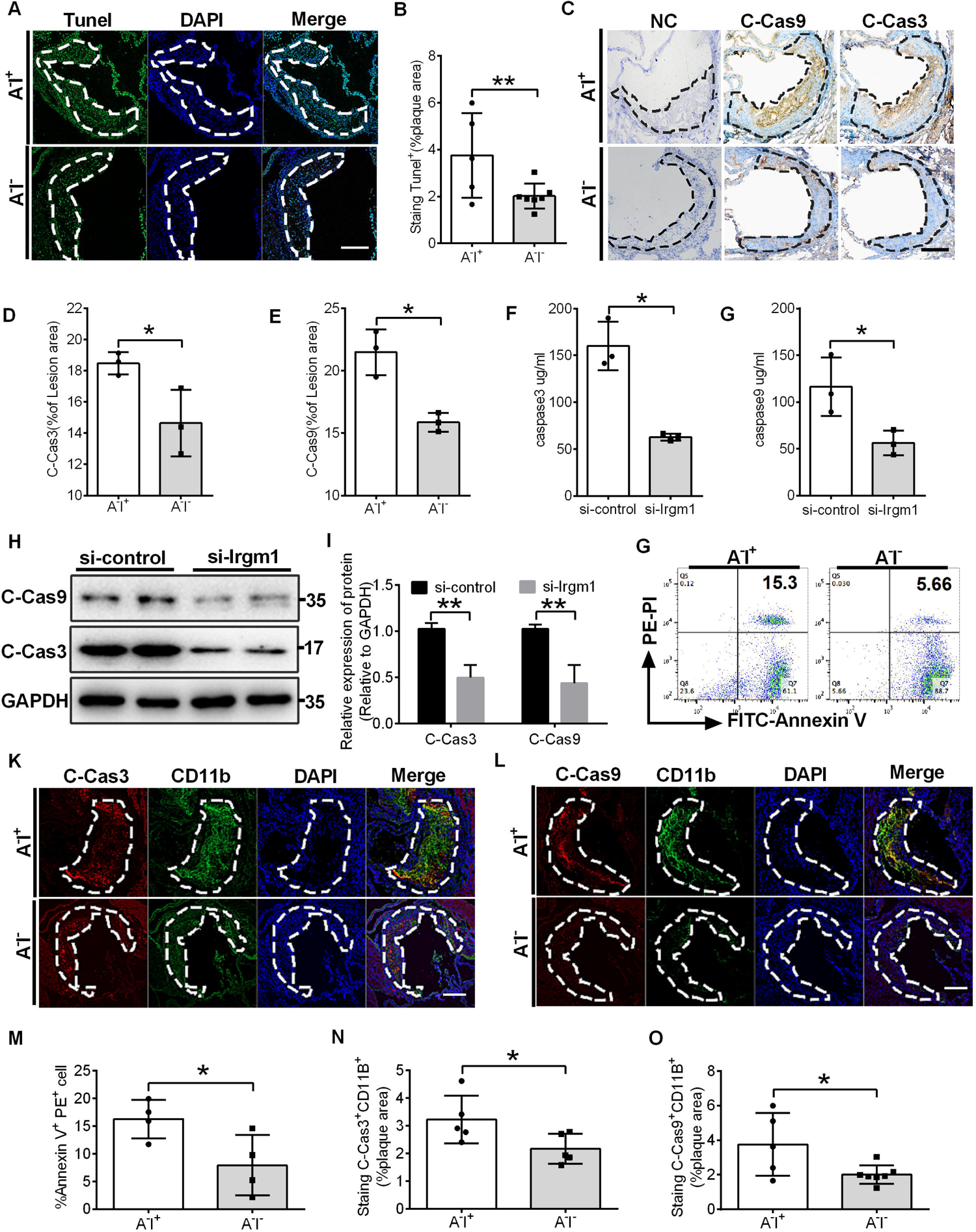
Irgm1 induces apoptosis in macrophages. (A) TUNEL^+^ cell areas in aortic sinus plaques from ApoE^-/-^Irgm1^+/+^ (n = 3) and ApoE^-/-^Irgm1^+/-^ mice (n = 3). Areas circled by dashed lines represent the contour of plaques; scale bars = 200 μm. (B) Quantification of the percentage of TUNEL^+^ areas within the plaques. (C-E) Representative images(C) and quantitative analyses (D, E) of cleaved-caspase3 and cleaved-caspase9 in the plaques (n = 3); NC=negative control; scale bars = 100 μm. (F–H) Raw264.7 cells were transfected with si-Irgm1, and then stimulated with ox-LDL (50 ng/ml) for 48 h. Activity of cleaved-caspase3 (F) and cleaved-caspase9 (G) was quantified. (H-I) Western blotting was used to detect the expression of cleaved-caspase3 and cleaved-caspase9(H) and quantitative analyses results were shown (I). Duplicate samples were run in the same gels, with the membrane cut in half. GAPDH was used as a loading control. Quantitative data represent the fold change after normalized to GAPDH. (J-O) ApoE^-/-^Irgm1^+/+^ and ApoE^-/-^Irgm1^+/-^ mice were given a high-fat diet for 16 weeks. (J, M) Peritoneal macrophages were recruited and then FITC-Annexin V and PE-PI were co-stained and quantified by flow cytometry (n = 4 in each group). (K) Representative images for co-location of cleaved-caspase3 (red) and CD11b (green) by immunofluorescence staining (n = 3 in each group). (L) Co-location of cleaved-caspase9 (red) and CD11b (green) by immunofluorescence staining (n = 3 in each group); nuclei stained by DAPI (blue); scale bars = 200 μ. (N, O) Quantitative data of the co-location percentage of CD11b^+^ with cleaved-caspase3 (N) and cleaved-caspase9 (O). A^-^I^+^= ApoE^-/-^Irgm1^+/+^ mice; A^-^I^-^= ApoE^-/-^Irgm1^+/-^ mice, \**p* < 0.05, \*\**P* < 0.01. Results are presented as the mean ± SD. Statistical analysis: unpaired Student’s *t-test*.

### ROS contributes to Irgm1-induced apoptosis in macrophages

Production and accumulation of ROS in macrophages contributes to apoptosis (28). We used DHE fluorescent probes to label ROS production in Raw264.7 cells. Compared with the si-control group, DHE fluorescence intensity of the si-Irgm1 group declined sharply (Figure 4A-4B), consistent with flow cytometry results (Figure 4C-4D). These results indicate that Irgm1 can promote production of intracellular ROS in macrophages. We next assessed whether ROS were involved in *Irgm1-*induced apoptosis. Compared with the si-control group, the expression of cleaved-caspase3 and cleaved-caspase9 were reduced in the si-Irgm1 group. N-acetylcysteine (NAC, an active oxygen scavenger) further reduced expression of cleaved caspase3 and caspase9 (Figure 4E-4F), and the activities of caspase3 and caspase9 showed similar results (Figure 4G–4H). Compared with the si-control group, the co-localized fluorescence intensity of TUNEL and DAPI in the si-Irgm1 group was significantly reduced which was, further decreased in the presence of NAC (Figure 4I–4J). In summary, Irgm1 induces apoptosis in macrophages by promoting production of ROS.

**Figure 4.**
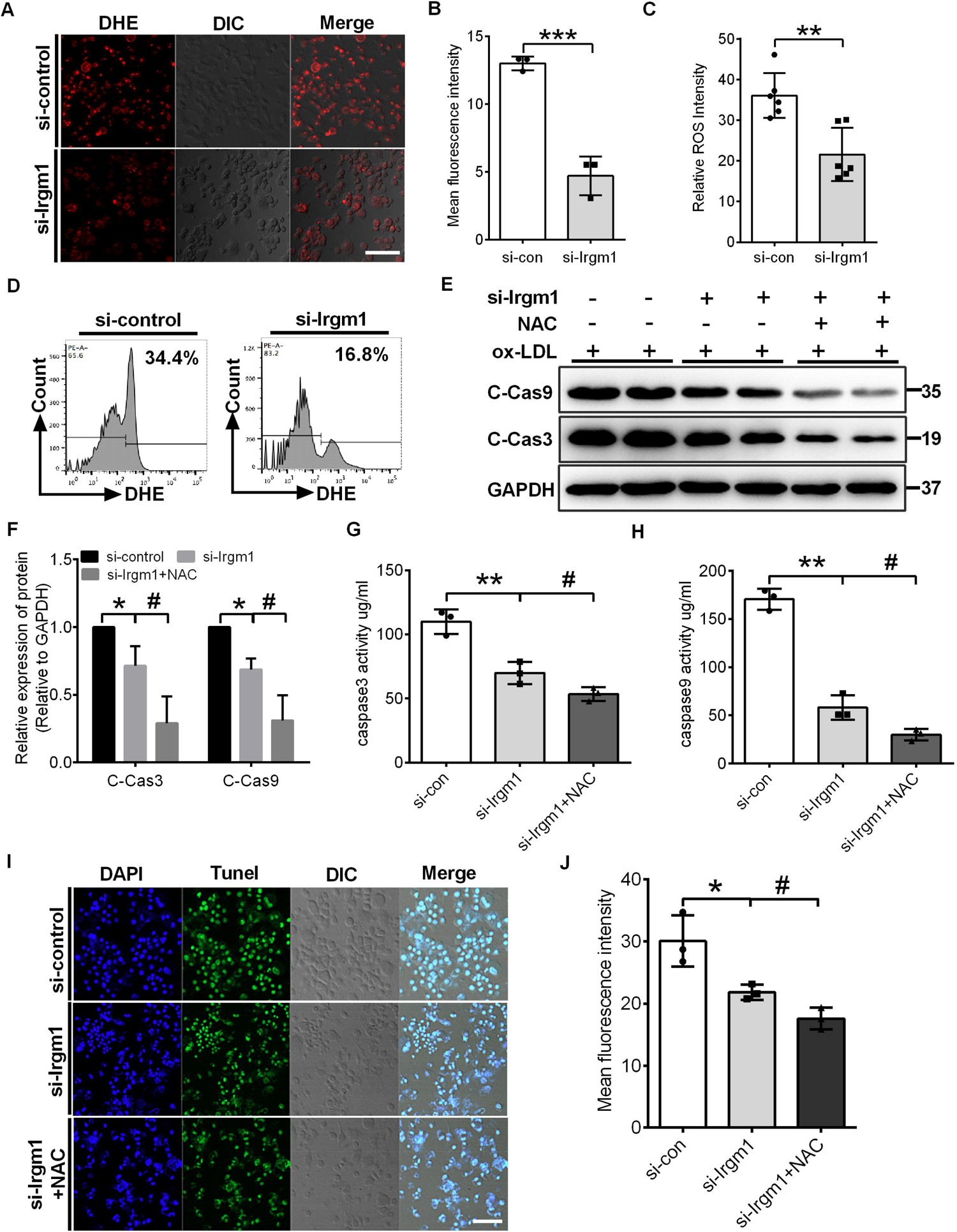
ROS contributes to Irgm1-induced apoptosis in macrophages. (A-D) The Raw264.7 cells were transfected with si-Irgm1 and si-control, and then μg/ml) for 48 h. (A) Representative images of the reactive oxygen species labeled with DHE (5 μM) fluorescent probe were observed by confocal microscopy (n = 3 per group); scale bars = 50 μ. (B) Quantitative data in the graph represent relative mean fluorescence intensity (MFI) (n = 3 per group). (C-D). Flow cytometry was used to detect ROS labeled with DHE fluorescent probes (n = 6 per group). (C) Quantitative data represent the percentage of DHE^+^ macrophages and representative graphs showing dot plots (D). (E-J) Raw264.7 cells transfected with si-Irgm1 and si-control, in presence or absence of NAC, and then stimulated with ox-LDL (50 μg/ml) for 48 h. (E) Western blotting was used to detect the expression of cleaved-caspase3 and cleaved-caspase9. Duplicate samples were run in the same gels, with the membrane cut in half. GAPDH was used as a loading control. (F) Quantitative data represent the fold change after normalization to GAPDH. (G, H) The protein activity of cleaved-caspase3 and cleaved-caspase9 was detected by ultraviolet spectrophotometry (n = 3 in each group). (I-J) Representative images(I) and quantitative data(J) for TUNEL^+^ macrophages (green) by immunofluorescence. nuclei were stained by DAPI (blue). DIC channel shown the contour of cells; scale bars = 50 μm; (n = 3 per group). si-control vs. si-Irgm1, \**p* < 0.05, \*\**p* < 0.01; si-Irgm1 vs. si-Irgm1+NAC, #*p* < 0.05, ##*p* < 0.01. Results are presented as the mean ± SD. Statistical analysis: unpaired Student’s *t-test*.

### Irgm1 induces apoptosis in macrophages by activating MAPK signaling pathway

MAPK signaling has been implicated in apoptosis through the JNK/P38/ERK pathway in response to various stress stimuli, including ROS (29). To explore how Irgm1 leads to macrophage apoptosis, we detected phosphorylated (p)-JNK, p-p38, and p-ERK in the MAPK signaling pathways, as well as the downstream apoptosis-related proteins Bax, Bcl2, cleaved-caspase3, and cleaved-caspase9. Compared with the si-control group, expression of p-JNK, p-p38, p-ERK, Bax, cleaved-caspase3, and cleaved-caspase9 were significantly decreased in the si-Irgm1 group. Bcl2 expression was increased (Figure 5A Lane 1,2 vs. Lane 3,4; Figure 5B–5C). NAC further decreased expression of p-JNK, p-p38, p-ERK, and downstream apoptotic proteins Bax, cleaved-caspase3, and cleaved-caspase9 in the si-Irgm1 group and increased expression of Bcl2 (Figure 5A Lane 5,6; Figure, 5B–5C). These results suggest that activation of the MAPK signaling pathway and ROS generation were major characteristics of Irgm1-induced apoptosis. To test this conclusion, the macrophages were treated with three inhibitors, including SP600125 (SP), SB203580 (SB) and U0126 (U), which inhibit JNK, P38, and ERK, respectively. Expression of JNK, P38, and ERK was significantly reduced by these inhibitors (Supplementary Figure 4A–F). In both si-control and si-Irgm1 groups, ox-LDL significantly increased expression of Bax, cleaved-caspase3, and cleaved-caspase9, and decreased expression of Bcl2 (Figure 5D Lane 1 vs. 2; Lane 6 vs. 7; Figure 5E–H). Importantly these inhibitors prevented the changes of MAPK signaling, pathway and apoptosis-related proteins (Figure 5D, lane 3–5 vs. lane 2; Lane 8–10 vs. lane 7; Figure 5H). Together, these results show that Irgm1 induces ROS production, which activates the MAPK signaling pathways, leading to apoptosis in macrophages.

**Figure 5.**
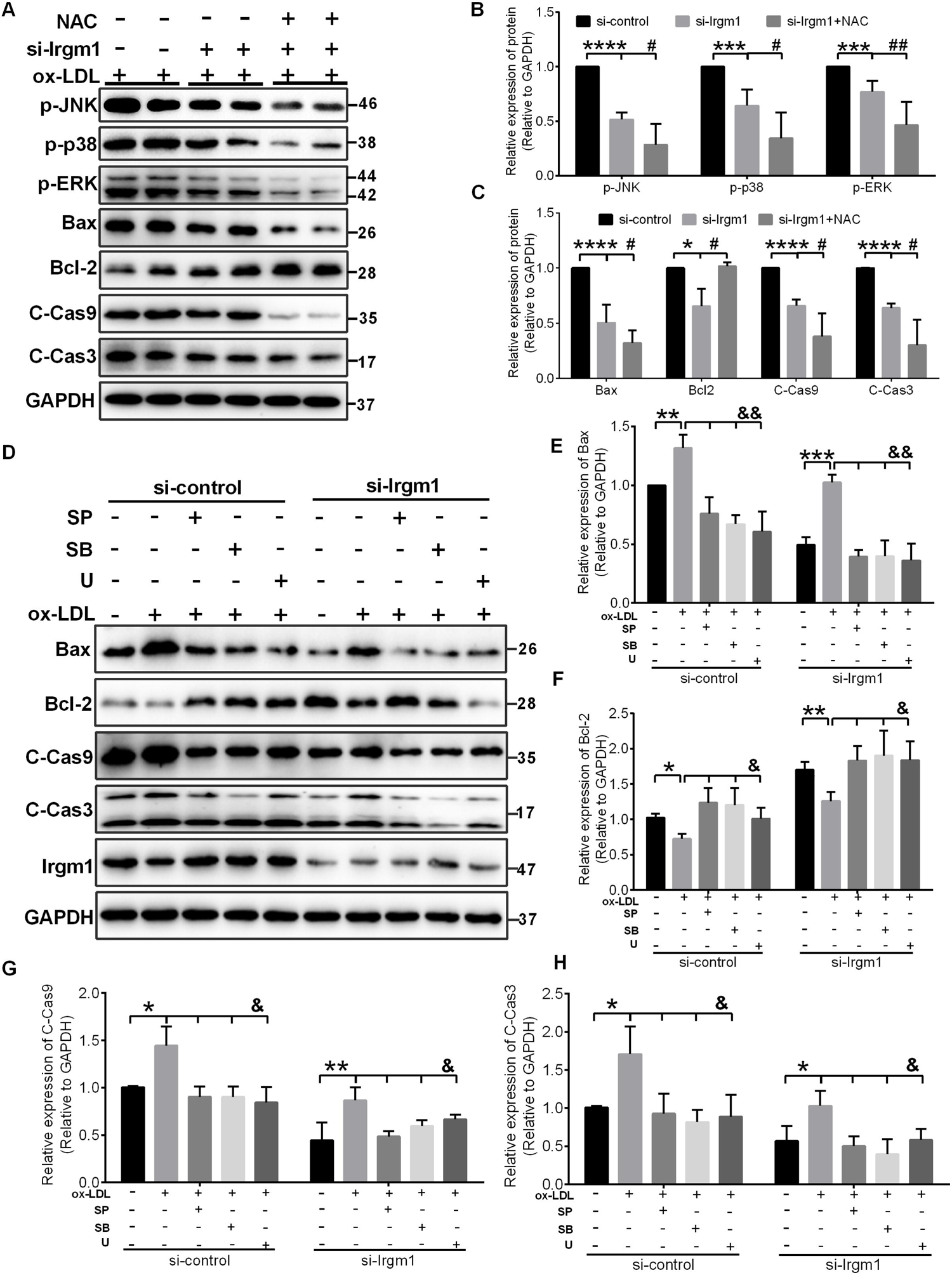
Irgm1 induces apoptosis in macrophages by activating the MAPK signaling pathways. (A-C) The Raw264.7 cells were transfected with si-Irgm1 and si-control, in presence or absence of NAC, and then stimulated with ox-LDL (50 μg/ml) for 48 h. Representative image(A) and quantitative data(B-C) of p-JNK, p-38, p-ERK and apoptosis-related Bax, Bcl2, cleaved-caspase3 and cleaved-caspase9 protein levels. GAPDH was used as a loading control. Quantitative data representing the fold change after normalized to GAPDH. (D) Expression of Bax, Bcl2, cleaved-caspase3 and cleaved-caspase9 were detected by western blotting in si-control (Lane 1–5) and si-Irgm1 (Lane 6–10) groups, after treatment with SP600125 (10 μ), SB203580 (10 μ) or U0126 (10 μ) for 3 h. GAPDH was used as a loading control. (E–H) Quantitative data representing the fold change after normalization to GAPDH. si-control vs. si-Irgm1 in (B, C): \**p* < 0.05, \*\**p* < 0.01, \*\*\**p* < 0.001, \*\*\*\**p* < 0.0001; si-Irgm1 vs. si-Irgm1+NAC, #*p* < 0.05, ##*p* < 0.01; si-control vs. si-Irgm1 in (D–H): Lane 1 vs. Lane 2, Lane 6 vs. Lane 7, \**p* < 0.05, \*\**p* < 0.01, \*\*\**p* < 0.001; Lane 3, 4, 5 vs. Lane 2; Lane 8, 9, 10 vs. Lane 7, &*p* < 0.05, &&*p* < 0.01. Results are presented as the mean ± SD. Statistical analysis: unpaired Student’s *t-test*.

### Irgm1 aggravates vulnerability of plaque and macrophage apoptosis in bone marrow chimera mice

We constructed chimera mice to investigate whether Irgm1 expressed on hematopoietic origin cells contributes to atherosclerosis progression. Bone marrow macrophages from ApoE^-/-^Irgm1^+/+^ or ApoE^-/-^Irgm1^+/-^ mice were transferred to ApoE^-/-^Irgm1^+/-^ mice (Figure 6A). Compared with ApoE^-/-^Irgm1^+/+^ mice, cholesterol crystals, thickness of fibrous cap, necrotic core area, ratio, and plaque area were significantly reduced in plaques of ApoE^-/-^Irgm1^+/-^ mice (Figure 6B–6E, 6H, Supplemental Figure 5A). ORO staining showed that the ratio of neutral lipids in ApoE^-/-^Irgm1^+/-^ mice plaques was reduced (Figure 6F, 6I). Compared with ApoE^-/-^Irgm1^+/+^ mice, both content and ratio of collagen fibers in plaques of ApoE^-/-^Irgm1^+/-^ mice were significantly reduced (Figure 6G, 6J; Supplemental Figure 5C). These results were consistent with those in non-recipient mice (Figure 3).

**Figure 6.**
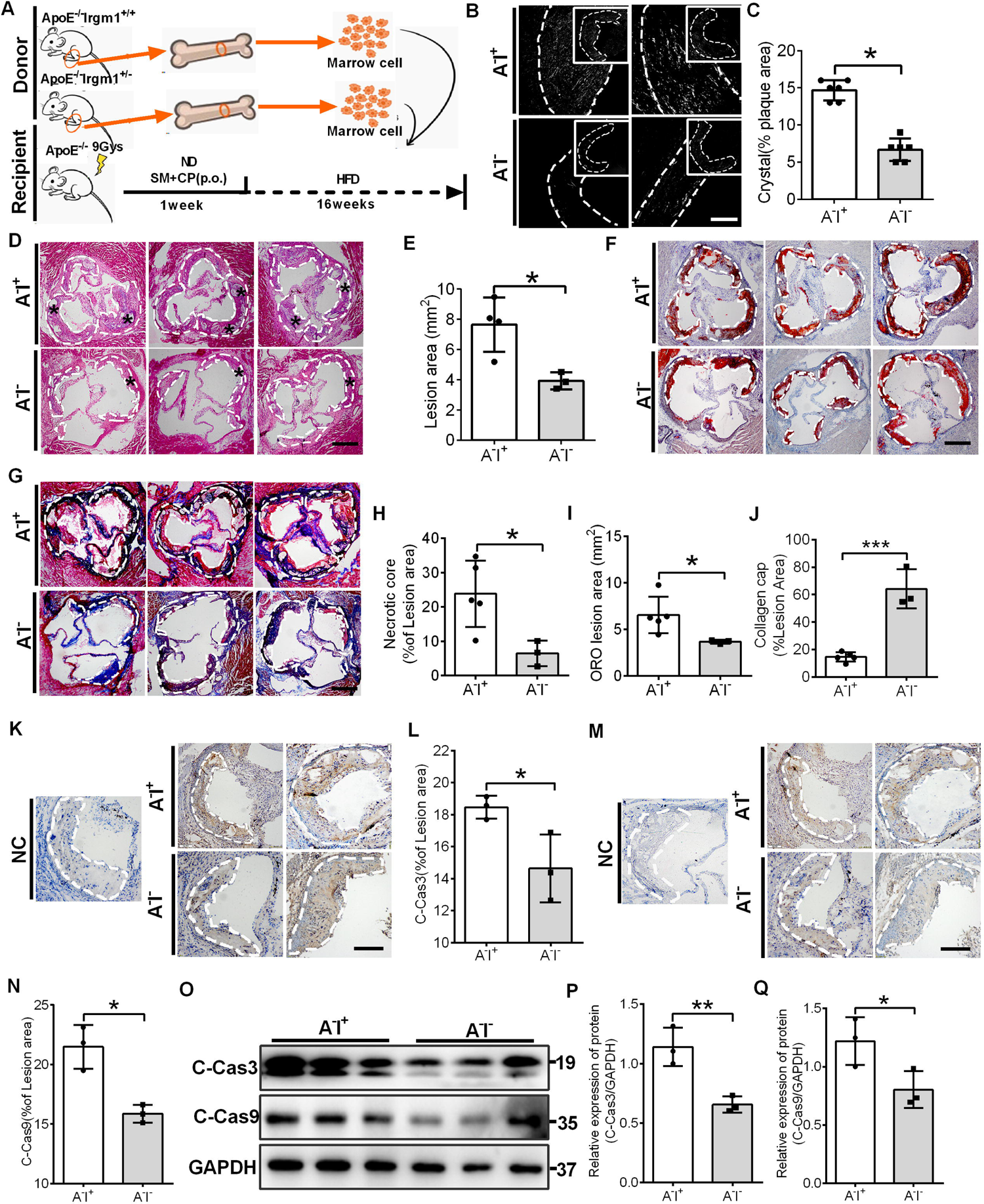
Irgm1 aggravates plaque progression in bone marrow chimera recipient mice and contributes to macrophage apoptosis. (A) Schematic diagram for the establishment of the bone marrow chimera recipient mouse model. Representative images (B) and quantitative analyses (C) of cholesterol crystals from ApoE^-/-^Irgm1^+/+^ (n = 3) and ApoE^-/-^Irgm1^+/-^ (n = 3) bone marrow chimera recipient mice; scale bars = 100 μ (D) Representative images for H&E staining of plaques in the aortic sinus from ApoE^-/-^Irgm1^+/+^ (n = 5) and ApoE^-/-^Irgm1^+/-^ (n = 3) bone marrow chimera recipient mice. A necrotic lipid core is indicated by *; dashed lines indicate the contour of the plaques; scale bars = 200 μm. (E, H) Quantitative analyses of FCT, and percentage of necrotic core areas in the plaques. (F) Representative images of neutral lipid in the aortic sinus detected by ORO staining from ApoE^-/-^Irgm1^+/+^ (n = 5) and ApoE^-/-^Irgm1^+/-^ (n = 3) bone marrow chimera recipient mice; scale bars = 200μm. (G) Representative images of collagen in the aortic sinus detected by Masson staining from ApoE^-/-^Irgm1^+/+^ (n = 5) and ApoE^-/-^Irgm1^+/-^ (n = 3) bone marrow chimera recipient mice; scale bars = 200 μ. (I, J) Quantitative analyses of the percentage of neutral lipid and collagen content in the plaques, respectively. (K-L) Representative images(K) and quantitative analyze(L) of cleaved-caspase3 in aortic sinus plaques. NC=negative control; scale bars = 100 μm. (M) Western blotting was used to detect the cleaved-caspase3 and cleaved-caspase9 protein levels of the aortic arch from ApoE^-/-^Irgm1^+/+^ (n = 5) and ApoE^-/-^Irgm1^+/-^ (n = 3) bone marrow chimera recipient mice. GAPDH was used as a loading control. (N) Quantitative data represent the fold change after normalized to GAPDH. ApoE^-/-^Irgm1^+/+^ vs. ApoE^-/-^Irgm1^+/-^, \**p* < 0.05, \*\**p* < 0.01. Results are presented as the mean ± SD. Statistical analysis: unpaired Student’s *t-test*.

To further test the contribution of Irgm1-induced apoptosis in chimera mice macrophages, we examined expression of cleaved-caspase3 and cleaved-caspase9 in the aortic sinus. Compared with ApoE^-/-^Irgm1^+/+^ mice, the ratio of cleaved-caspase3 and cleaved-caspase9 in plaques of ApoE^-/-^Irgm1^+/-^ mice was significantly reduced (Figure 6K–6N), consistent with western blotting results (Figure 6O–6Q). Finally, we observed co-localization of cleaved-caspase3 and cleaved-caspase9 with CD68 (Supplemental Figure 5D–5E). In summary, Irgm1 aggravates vulnerability of plaque and contributes to apoptosis in macrophages in chimera mice.

## Discussion

In the STEMI cohort, serum IRGM levels were increased in patients with culprit PR (Figure 1B). This potentially provides a tool to predict PR. When adjusted for other risk factors related to ACS, IRGM was still correlated with PR (Figure 1C). Apparently, IRGM/Irgm1 affects the vulnerability of atherosclerotic plaques. In this study, we clarified how Irgm1 affects plaque phenotype *in vivo* and *in vitro*. Irgm1 activates the MAPK signaling pathway by promoting ROS production, leading to macrophage apoptosis and, thus, affecting plaque stability (Figure 7). These data identify a crucial role for Irgm1 in mediating apoptosis of macrophages during the progression of atherosclerosis, and highlight IRGM as a potential predictor of atherosclerotic plaque vulnerability.

**Figure 7.**
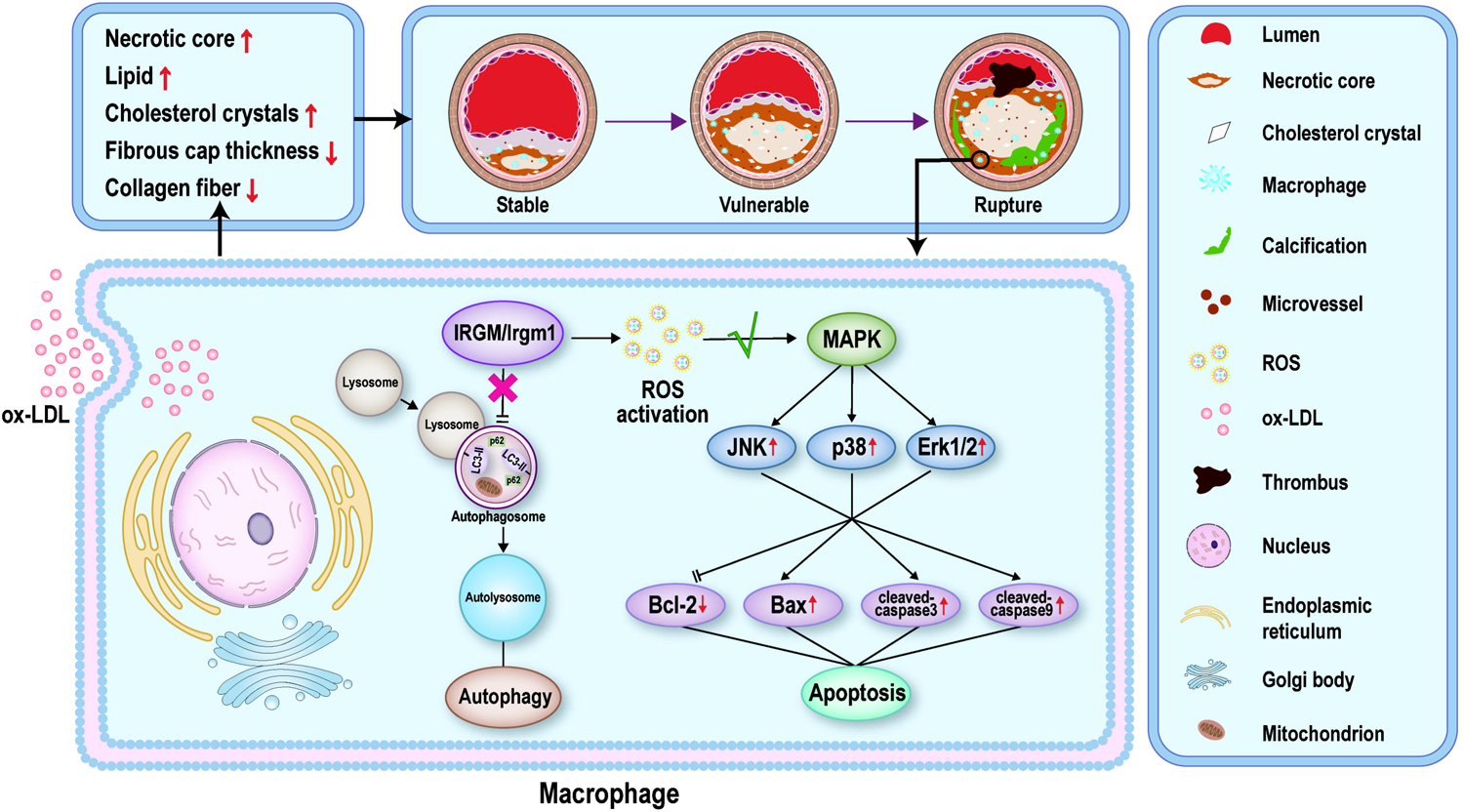
Summary of the role of IRGM/Irgm1 in atherosclerotic plaque progression by promoting macrophage apoptosis. During the progression of atherosclerotic plaque, ox-LDL is continuously engulfed by the increasing number of macrophages. The expression of IRGM/Irgm1 in macrophages increases rapidly, which promotes the production of ROS, activates the MAPK signaling pathway, and finally induces macrophage apoptosis. The stability of the plaque then changes, with increases in the content of necrotic core, lipid, and cholesterol crystals, fibrous cap thinning, and a decrease in collagen fibers, transforming the plaque from stable to vulnerable. A vicious circle leads to further plaque progression and eventual plaque rupture.

In the 1980s, Davies *et al*. showed that PR caused most fatal myocardial infarctions (32). It became accepted that instability of coronary atherosclerosis is due to weakening of the collagen structure caused by inflammatory infiltration and fissure of TCFA plaques (33–34). The emergence of OCT makes it possible to track the pathogenesis of ACS in patients *in vivo* and assess the morphological features of vulnerable plaques (35). Our previous studies found that most of the plaques that tend to rupture have thin fibrous caps and large lipid arcs, as assessed by OCT combined with intravascular ultrasound (7). Here, we compared the morphological characteristics between ruptured and non-ruptured culprit plaques in patients with STEMI. We confirmed that the fibrous cap of the ruptured plaque was thinner and contained more lipids (Figure 1A). However, there is no good method for the early diagnosis and treatment of PR, mandating a search for biological predictors of PR in ACS patients.

Inflammation plays essential roles in the progression from vulnerable to ruptured plaque, ultimately leading to clinical events (36). Administration of IL-1β monoclonal antibody provided a 15% reduction in the risk of further MI, stroke, or death, proving the effectiveness of anti-inflammatory treatment on atherosclerotic diseases (37). Hunn *et al*. have shown that IRGM belongs to the family of interferon-induced GTPases (IRGs), genetically and functionally related to proteins implicated in chronic inflammatory diseases (38–39). Therefore, we analyzed expression of serum IRGM in patients with STEMI and found that serum IRGM was significantly increased in the RUP group (Figure 1B). Considering TNF-α and IL-6 play central roles in α downstream inflammatory responses that lead to atherosclerosis (40), we quantified expression of serum IRGM, IL-6, and TNF-α, and found that expression of IRGM α was positively correlated with IL-6 and TNF-α (Supplemental Figure 2D–G). These α findings indicate that IRGM/Irgm1 plays a potential role in the progression from vulnerable to ruptured plaque, ultimately leading to clinical events.

IRGM/Irgm1 has been extensively studied in inflammation and cancer (41, 42). IRGM/Irgm1 promotes macrophage autophagy and protects the intestine from inflammatory attack (39, 43). Human IRGM proteins are targeted by RNA viruses, some of which improve their own replication ability by autophagy (44, 45). Inhibiting the expression of IRGM/Irgm1 by microRNA attenuates autophagy by macrophages (46). Here, we confirmed that IRGM was highly expressed in the RUP group, which was related to the vulnerability of advanced atherosclerotic plaques (Figure 1). We next investigated whether the IRGM/Irgm1-induced macrophage autophagy contributes to vulnerability of atherosclerotic plaques. After a 3-h treatment with ox-LDL, Irgm1 up-regulated mRNA and protein levels of autophagy indicators LC3, P62, Atg5 and beclin1. Treatment with si-Irgm1 increased RFP-positive/GFP-positive puncta, suggesting that si-Irgm1 inhibits autophagosome–lysosome fusion (Supplemental Figure 1). Unfortunately, after stimulating RAW264.7 cells with ox-LDL for 48 h, only RFP-positive/GFP-positive puncta were observed in both the si-Irgm1 and si-control groups. These results suggested that in the early stage of atherosclerosis, Irgm1 may prevent induction of autophagy in macrophages, whereas the autophagy flux of macrophages is blocked in an advanced atherosclerotic model (Supplemental Figure 1). Therefore, the effect of IRGM on macrophage autophagy cannot explain changes in plaque vulnerability in advanced atherosclerosis.

Evidence has amassed that macrophage apoptosis intensifies formation of plaque necrotic cores, causing thinning of the fibrous cap, and ultimately leading to PR (25, 47) (48–50). In this study, we first explored the effect of Irgm1 on atherosclerotic plaque phenotype in mice. The aortic plaque area, cholesterol crystals, necrotic core, and neutral lipids were reduced in ApoE^-/-^Irgm1^+/-^ mice, whereas collagen fibers increased. These results indicate that Irgm1 can aggravate the vulnerability characteristics of advanced atherosclerotic plaques (Figure 2). Next, we found that Irgm1 promotes expression of apoptosis-related proteins cleaved-caspase3 and cleaved-caspase9 in macrophages (Figure 3), also confirmed *in vitro*. Finally, Irgm1 deficiency in bone marrow *chimera* mice displayed significant delays in atherosclerotic progression and had stabilized plaques (Figure 6). In summary, IRGM/Irgm1 contributes to plaque vulnerability by promoting macrophage apoptosis in advanced atherosclerotic plaques but mechanisms remain unclear. Hypotheses proposed to explain the apoptosis of macrophages in advanced atherosclerosis include deprivation of growth factors or the presence of toxic cytokines (51, 52), yet there is no evidence for these hypotheses *in vivo*.

Many studies have explored the continuous accumulation of ROS, triggering depolarization of the inner mitochondrial membrane and eventually leading to cell apoptosis (48, 59). Markers of endoplasmic reticulum stress and macrophage apoptosis are found in human atherosclerotic plaques. Here, we detected reactive oxygen fluorescent probes *in vitro* and found that Irgm1 can promote ROS production in macrophages. Interestingly, ROS scavenger NAC and si-Irgm synergistically reduce levels of apoptosis-related proteolytic caspase3 and caspase9 in RAW264.7 cells. These results suggest that Irgm1 can lead to apoptosis *in vitro* by promoting ROS production in macrophages. Increased accumulation of ROS activates MAPK (53–56). Other studies have shown that the MAPK signaling pathway is regulated by ERK1/2, c-Jun N-terminal kinase1/2/3 (JNK1/2/3), and p38-MAPK in the processes of autophagy, apoptosis, and inflammation (61, 62). We speculate that Irgm1 promotes ROS production in macrophages and activates apoptosis via the MAPK signaling pathway. Compared with the control group, when macrophages were treated with ERK1/2, JNK, and p38-MAPK inhibitors, expression of the apoptotic proteins BAX, Bcl-2, *cleaved-caspase*3, and *cleaved-caspase*9 was significantly reduced (Figure 5). In summary, these results indicate that Irgm1 can activate the MAPK signaling pathway and cause macrophage apoptosis.

These results are subject to limitations. First, as a retrospective study, the clinical data reflect the current epidemiological, serum, and plaque features of Chinese patients with STEMI. It may not be generalizable to patients with NSTE-ACS or patients from other locations. Second, although macrophage infiltration is defined in a published consensus on OCT, it remains a controversial topic to recognize macrophage infiltration by OCT. Therefore, this study did not list the data of macrophage infiltration by OCT between the RUP and NO-RUP groups. Third, as we could not obtain coronary arteries from clinical patients, our research group used lower limb arteries obtained from vascular surgery to assess plaque characteristics. This may not accurately reflect the profile of coronary plaques. Fourth, since ApoE^-/-^Irgm1^-/-^ mice in the C57BL/6 background may be embryonic lethal, we have been unable to obtain the above double-gene knockout mice. Here, we were only able to use ApoE^-/-^Irgm1^+/-^ mice as the experimental group *in vivo*. Fifth, it may be important to study the progression of atherosclerotic plaques by establishing an atherosclerosis model through a high-fat diet. However, this modeling method may not be representative for the study of PR. We are actively seeking a way to construct a PR model to compensate for drawbacks of this research. Based on our conclusions, we have abundant reasons to regard Irgm1 as a suitable predictive factor for rupture in the rupture model of advanced atherosclerotic mice. Sixth, we found that Irgm1 regulates macrophage apoptosis by activating the MAPK signaling pathway. Others and we have not yet verified the specific site at which Irgm1 acts in regulating the MAPK signaling pathway. Seventh, in the sequence of IRGM (+313) (rs10065372), an SNP is present at position +313 inside exon 2 of IRGM at chromosome 5q33.1. This SNP is associated with severe sepsis with a high fatality rate. It is not clear whether mouse Irgm1 has the same SNP as human IRGM. We must keep in mind these limitations, but our findings reinforce each other due to the multiplicity of approaches.

## Methods

### Study Participants

In this study, 135 patients who were diagnosed with STEMI in the 2nd Affiliated Hospital of Harbin Medical University (Harbin, China) between August and December 2016 and who underwent OCT after thrombus aspiration were enrolled. Main exclusion criteria were angina pectoris, myocarditis, heart failure, aortic dissection, acute and chronic infections and infectious diseases, liver and kidney insufficiency, and severe progressive diseases. We finally enrolled 85 patients with STEMI. We divided patients into two groups: patients with PR (RUP group) (n = 52) and without PR (NO-RUP group) (n = 33). Eight additional healthy volunteers were also recruited. We analyzed OCT images and obtained the serum of these 85 patients and 8 healthy volunteers. The definition of STEMI was in accordance with the 2015 ACC/AHA/SCAI guidelines published previously (63).

### OCT Image Acquisition and analysis

The frequency-domain OCT ILUMIEN system, along with the Dragon fly catheter (St. Jude Medical, Westford, Massachusetts, USA) were used to acquire OCT imaging of culprit lesions, as shown previously (9). Two experienced independent investigators (H.C and S.S) analyzed OCT images according to the criteria of the Consensus OCT Document (64) using a proprietary OCT review workstation (St. Jude Medical, Westford, MA, USA) at an intravascular imaging and physiology core lab. If these two observers disagreed, a consensus reading was obtained by judgement with another investigator. Investigators were blinded to the subject information. Plaque rupture was defined as a lipid plaque with fibrous cap continuity destroyed plus a clear cavity formation within the plaque. The FCT covering the lipid core was measured three times at the thinnest location, and the mean value of triplicate measurements was calculated. TCFA was a lipid plaque in the presence of FCT < 65 μm with a maximum lipid arc of at least 90° (65).

### Patient blood vessel specimens

Atherosclerotic middle artery vessels were obtained from patients with arterial occlusion in the Department of Vascular Surgery of the First Affiliated Hospital of Harbin Medical University. Normal vessels were obtained from five patients who underwent lower limb amputation. Blood vessel specimens from patients were transported to the laboratory by dry ice and maintained at low temperature (−40 °C) after isolation. Blood vessel specimens were embedded in paraffin and sectioned consecutively (4 μm) for H&E and immunohistochemical staining.

### Animals

Irgm1^-/-^ mice (C57BL/6 background) were derived from a published source (17). Irgm1^-/-^ mice and ApoE^-/-^ mice were crossed to obtain ApoE^+/-^Irgm1^+/-^ mice. We screened ApoE^-/-^Irgm1^+/-^ mice by crossing ApoE^+/-^Irgm1^+/-^ mice. Due to the probability of embryonic lethality in ApoE^-/-^Irgm1^-/-^ mice, we genetically identified offspring of self-crossed ApoE^-/-^Irgm1^+/-^ mice. After PCR genotyping, we then established a model of advanced atherosclerosis in mice fed high-fat diet for sixteen weeks.

### Reagents

See Supplemental Table 3–4.

### Bone marrow chimera recipient mice

ApoE^-/-^Irgm1^+/+^ (6–8 weeks old) recipient mice were supplemented with fradiomycin, polymyxin, and sterile pH 3.0 water for a week. They were then irradiated with 9 Gy of radiation.

The recipient mice were divided into three groups. Two groups of recipient mice received bone marrow macrophages (5**Error! Reference source not found.**10^6^) from ApoE^-/-^Irgm1^+/+^ donor mice and ApoE^-/-^Irgm1^+/-^ donor mice by tail vein injection, whereas the control group received saline. After about seven days, the control group died and the experimental group survived, indicating that the modeling was successful.

### Cell culture

Raw264.7 cells were cultured in Dulbecco’s Modified Eagle’s medium (DMEM) (SH30022.01B, HYCLONE, USA) containing 10% FBS (0500, ScienCell, USA). Cells were seeded in 60 mm dishes or 6-and 12-well plates and collected at 60%–80% confluency.

### Transfection

Transfection was accomplished using Lipo3000 (L3000008, Invitrogen, USA) according to manufacturer’s protocols with the following siRNA duplexes: negative control, 5’-CGUACGCGGAAUACUUCGAUU-3’; Irgm1-siRNA, 5’-GGGCUGGGAUUCUGUCAUA-3’. Real-time qPCR was utilized to confirm efficiency of transfection.

### ROS detection

ROS generated by ox-LDL treatment in cells was evaluated by fluorescence intensity of the dihydroethidium (DHE) probe. After the intake of ox-LDL, macrophages were harvested, washed, and incubated with 5 μ DHE probe (S0063, Beyotime, China) for 30 min at 37 °C in the dark. After rinsing, fluorescent signals were immediately measured using a fluorescence microscope and FACS Verse flow cytometer (BD, USA).

### Real-time quantitative

*PCR (RT-qPCR).* Total RNA extracted by TRIzol reagent (Thermo Fisher, USA) was reverse-transcribed utilizing the RT Easy II First Strand cDNA Synthesis Kit (04379012001, Roche, Switzerland). We amplified cDNA (18 ng) in a Real-Time PCR Easy (SYBR Green I) (HY-K0501, MCE, China) on Bio-RAD Sequence Detection system (Bio-RAD, USA). The following primers were used in Supplemental Table 5, Gene expression values were normalized against β-actin.

### Pathological staining

We placed hearts in a mold containing tissue-freezing medium, which was perfused with PBS and frozen. Eight μm-thick (H&E staining) and twenty μm-thick (ORO staining) sections were cut from the hearts, from caudal of the aortic sinus to the proximal aorta. Slides were fixed with 4% paraformaldehyde and then stained. H&E, ORO, and Masson staining are as previously described (17).

### Immunofluorescence

Frozen slides were rewarmed at RT for 10 min, and then fixed with cold acetone (−20 °C) for 10 min. After rinsing three times in distilled water, slides were incubated with 0.3% Triton-X 100 for 30 min at 37 °C, and incubated with primary antibody at 4 °C overnight. On day 2, after rewarming for 10 min at room temperature (RT), slides were incubated with secondary antibody at RT for 1 h. Finally, nuclei were stained with 0.5 g/L DAPI for 10 min and images were captured using a confocal laser scanning microscope (ZEISS LSM 700).

### Immunohistochemistry

Frozen slides were rewarmed at RT for 10 min, and then fixed with cold acetone (−20 °C) for another 10 min. After three rinses in distilled water, slides were incubated with 0.3% Triton-X 100 for 30 min at 37 °C. Endogenous peroxidase was blocked with 3% hydrogen peroxide, and slides were incubated with primary antibody at 4 °C overnight. On the next day, frozen slides were rewarmed for 10 min at RT and then incubated with secondary antibody at RT for 1 h. Next, slides were exposed to 3,3’-diaminobenzidine (DAB) (ZSGQ-BIO, China) stain for 1–10 min, followed by Hematoxylin stain. Images were taken by ordinary forward microscopy (Olympus, BX41). Paraffin sections were first dewaxed and then fixed with 4% PFA for 5 min. After rinsing with PBS, the sections were incubated with 0.3% Triton-X 100 at 37 °C for 30 min, and then antigen repair proceeded. The rest of the steps were the same as for frozen sections.

### Western blotting

Nuclear and cytosolic fractions were obtained using a nuclear/cytosol fractionation kit (R0050, Solarbio, China). Total protein was extracted using RIPA buffer with phenylmethylsulfonyl fluoride, and Halt Protease and Phosphatase Inhibitor. Protein concentration was measured by a bicinchoninic acid protein assay. Protein samples (30 μg) were separated by sodium dodecyl sulfate-polyacrylamide gel electrophoresis (SDS-PAGE, P0015, Beyotime, China) and transferred to 0.22-μ m PVDF membranes. After blocking with 5% dried skimmed milk for 2 h at RT in Tris-buffered saline, the membranes were probed with primary antibodies (1:1000) and incubated overnight at 4 °C. Membranes were incubated for 2 h in the presence of horseradish peroxidase (HRP)-conjugated secondary antibodies (1:8000) at RT. Immunoreactivity was visualized by chemiluminescence using a ChemiDoc^TM^ MP Imaging System (Tanon, China). Protein bands were quantified using a Bio-Rad Chemi EQ densitometer and Bio-Rad Quantity One software (Tanon, China) and normalized to GAPDH.

### Caspase-3/9 Activity

Caspase-3/9 activity in the vascular tissue or cell homogenates was evaluated using the Caspase-3 Assay Kit or Caspase-9 Assay kit according to manufacturer’s instructions.

### Genotyping

Excised mouse toes were added to 100 μ (bimake) and digested for 15 min at 55 °C, then incubated for 5 min at 95 °C to inactivate enzymes. Tubes were centrifuged at 12,000 rpm for 5 min, with supernatant used as the PCR template (1000 ng). Primer sequences are listed in supplementary table 6. Next, PCR amplification was performed, followed by agarose gel electrophoresis and imaging analysis using a gel imaging system (BIO RAD, GelDoc Go).

### Autophagy with double-labeled adenovirus

Raw264.7 cells were transfected with si-Irgm1 for 24 h. Adenovirus vectors encoding LC3 (HBAD-mRFP-GFP-LC3, HANBIO, China) and ox-LDL (50 μg/mL) were added for 3 h or 48 h. Transfection efficiency was detected by confocal laser scanning microscope (ZEISS LSM 700). Autophagosomes were represented by the co-localized yellow fluorescence of both GFP and RFP. Due to quenching of GFP signal in acidic compartments, it shows stronger red fluorescence and less co-localization in autolysosomes. LC3 activation was represented by punctate dots in GFP-LC3 transfected cells.

### Detection of apoptotic cells by flow cytometry

Cells were cultured and incubated in a 5% CO2 atmosphere for 48 h at 37 °C and then harvested, washed twice with pre-chilled PBS, and resuspended in 200 µL of 1× binding buffer. The cells were stained with fluorescein isothiocyanate (FITC)-conjugated annexin V and propidium iodide (PI) using the Annexin V-FITC & PI Apoptosis Detection Kit (, BD 货号 Biosciences, USA).

### Statistics

Statistical analyses were performed using SPSS version 20.0 (SPSS, Inc., Chicago, IL, USA) or GraphPad Prism 7.0 (GraphPad Software). Continuous variables were expressed as mean ± SD for normally distributed variables and as median (25, 75th percentiles) for non-normally distributed variables, and assessed by Student *t-test* or Mann-Whitney U test, respectively. Categorical variables were presented as counts and percentages, and the comparisons were performed using a Chi-square test or Fisher’s exact-test. A series of multiple logistic regression models were used to assess the relationship between the risk of plaque rupture and plasma IRGM levels. A 2-sided *p* value < 0.05 was considered statistically significant. All data were analyzed by an independent statistician.

### Study approval

This study was approved by the Ethics Committee of the 2nd Affiliated Hospital of Harbin Medical University (Harbin, China), and all patients provided written informed consent. Animals were sacrificed by cervical dislocation. Animal experiments were performed according to guidelines approved by the Institutional Animal Care Committee and in compliance with the guidelines from Directive 2010/63/EU of the European Parliament with approval from the Harbin Medical University Ethics Review Board.

## Author contributions

SHF, SS, HXC, BS, and BY designed experiments. HXC and SS wrote the draft of the manuscript, followed by editing by SHF, YB, and JWT. SS, XRH, XYZ, JTT, ZYL, and ZZH performed the animal experiments; SHF, HXC, XW, SS, CCL, and WH performed experiments *in vitro*. ZMZ and CCL collected clinical data. HXC, SS, and JWT measured clinical images. SJW and HXC analyzed and visualized clinical results. BS and LMY provided crucial reagents. BY, BS, and JWT take responsibility for accuracy of the analysis of the whole experiment.

## Acknowledgments

Dr. Yu was supported by National Key R&D Program (No. 2016YFC1301100, 2016YFC1301101) and Major Instrument Development Project of the National Natural Science Foundation of China (No. 81827806). Dr. Fang was supported by National Natural Science Foundation of China (No. 81870353) and Natural Science Foundation of Heilongjiang Province (No. LH2020H048). The authors thank all participants and staff from the coronary interventional therapy team and the intravascular imaging core laboratory from the 2nd Affiliated Hospital of Harbin Medical University.

