## Supplementary file for "IRGM/Irgm1 Aggravates Progression of Atherosclerosis by Inducing Macrophage Apoptosis through the MAPK Signaling Pathway"

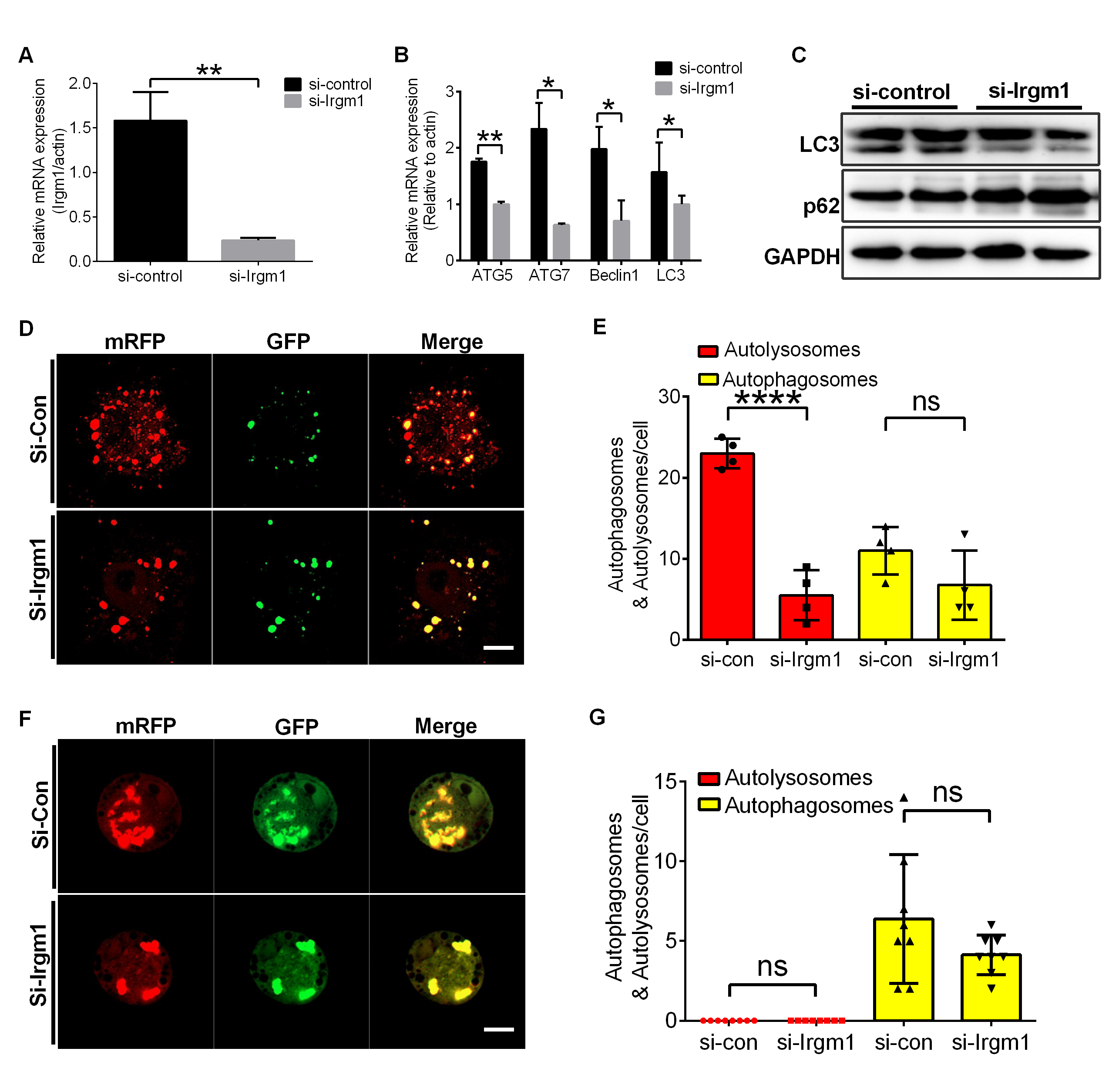

**Supplemental Figure 1**

(A) qPCR was used to detect the silencing efficiency in Raw264.7 macrophages after transfection with Irgm1 (n=3). (B) After stimulating Raw264.7 macrophages with ox-LDL (50 μg/ml) for 3 h, the expression of autophagy-related genes LC3, Atg5, Atg7, and Beclin1 was detected by qPCR (n=3). (C) Raw264.7 macrophages were stimulated with ox-LDL (50 μg/ml) for 3 h, after which the expression of autophagy-related proteins LC3 and P62 was detected by western blot. (D) Under the same treatment conditions as in (C), the autophagy double-labeled adenovirus fluorescent probe was used to detect the expression of autophagosomes (yellow) and autophagolysosomes (red). Green fluorescence was quenched in an acidic environment. Scale bar=10 μm. (E) Quantitative analysis of the yellow and red puncta in (D). (F) After stimulating Raw264.7 macrophages with ox-LDL (50 μg/ml) for 48 h, the autophagy double-labeled adenovirus fluorescent probe was used to detect the expression of autophagosomes (yellow) and autophagolysosomes (red). Green fluorescence is quenched in an acidic environment. Scale bar=10 μm. (G) Quantitative analysis of yellow and red puncta in (F). si-control vs. si-Irgm1, **p*<0.05, ***p*<0.01, *****p*<0.0001. Results are presented as mean ± SD. Statistical analysis: unpaired Student’s *t-test*.

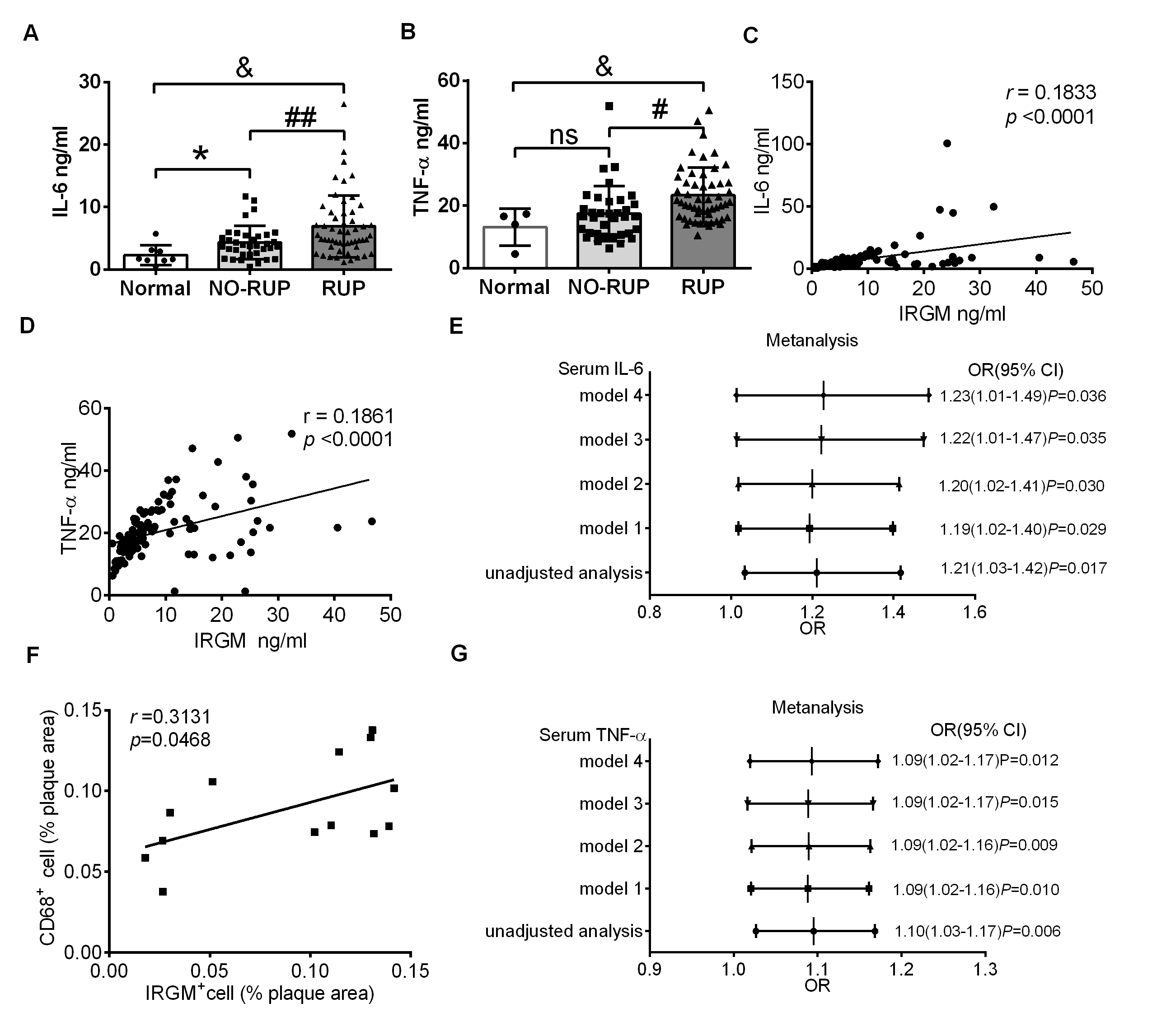

**Supplemental Figure 2**

(A) Serum IL-6 levels among healthy volunteers, NO-RUP patients, and RUP patients as detected by ELISA. **p*<0.05; ##*p*<0.01; &*p*<0.05. (B) Serum TNF-α levels among healthy volunteers, NO-RUP patients, and RUP patients as detected by ELISA. **p*<0.05; ##*p*<0.01; &*p*<0.05. (C) Quantitative data of the correlation between serum IRGM and IL-6 in patients with plaque rupture. (D) Quantitative data of the correlation between serum IRGM and TNF-α in patients with plaque rupture. (E, G) Adjusted ORs and 95% CIs for plaque rupture associated with serum TNF-α (E) and IL-6 (G) in patients with STEMI using 4 models: Model 1, a model adjusted for age; Model 2, in which Model 1 was further adjusted for hypertension, diabetes mellitus, and current smoking; Model 3, in which Model 2 was additionally adjusted for lipid factors (LDL-C and TC) and Model 4, in which Model 3 was further adjusted for peak TnI and hs-CRP. (F) Quantitative data of the correlation of IRGM^+^ and CD68^+^ area in the atherosclerotic plaque by immunohistochemistry.

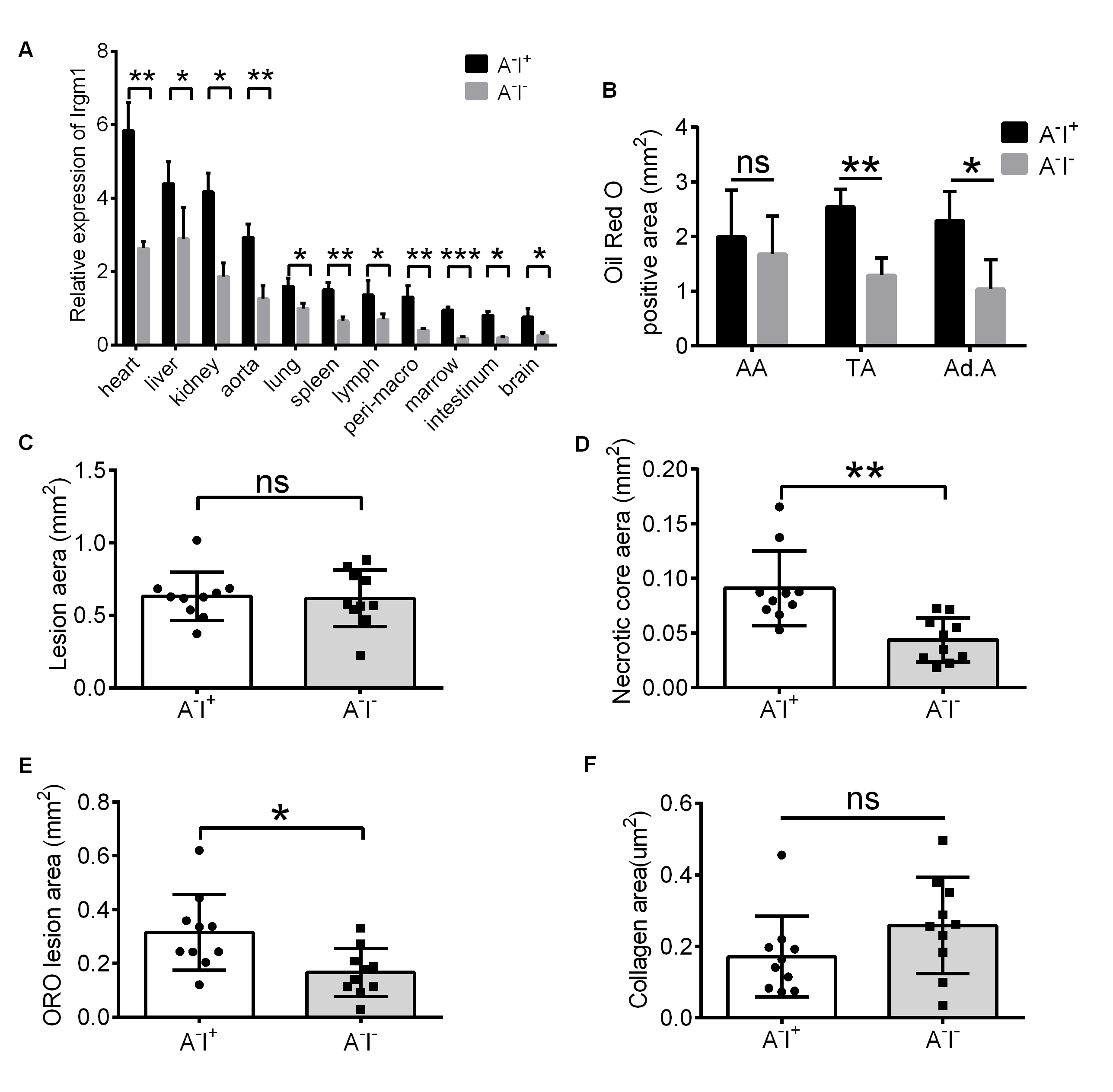

**Supplemental Figure 3**

(A) The expression of Irgm1 in the heart, liver, spleen, lung, kidney, aorta, intestine, brain, lymph nodes, peritoneal macrophages, and bone marrow macrophages in ApoE^-/-^Irgm1^+/+^ mice (n=3) and ApoE^-/-^Irgm1^+/-^ mice (n=3) as measured by qPCR. (B) Quantitative analysis of plaque area in the aortic arch, thoracic aorta, and abdominal aorta as determined by ImageJ software (n=4). (C–D) Quantitative analysis of the area of aortic sinus plaques (n=10) and necrotic cores (n=10) as determined by ImageJ. (E–F) Quantitative analysis of the area of neutral lipid (n=10) and collagen fiber (n=10) in aortic sinus plaques as determined by ImageJ. ApoE^-/-^Irgm1^+/+^ vs. ApoE^-/-^Irgm1^+/-^, **p*<0.05, ***p*<0.01, ****p*<0.001, ns=no significance. Results are presented as mean ± SD. Statistical analysis: unpaired Student’s *t-test*.

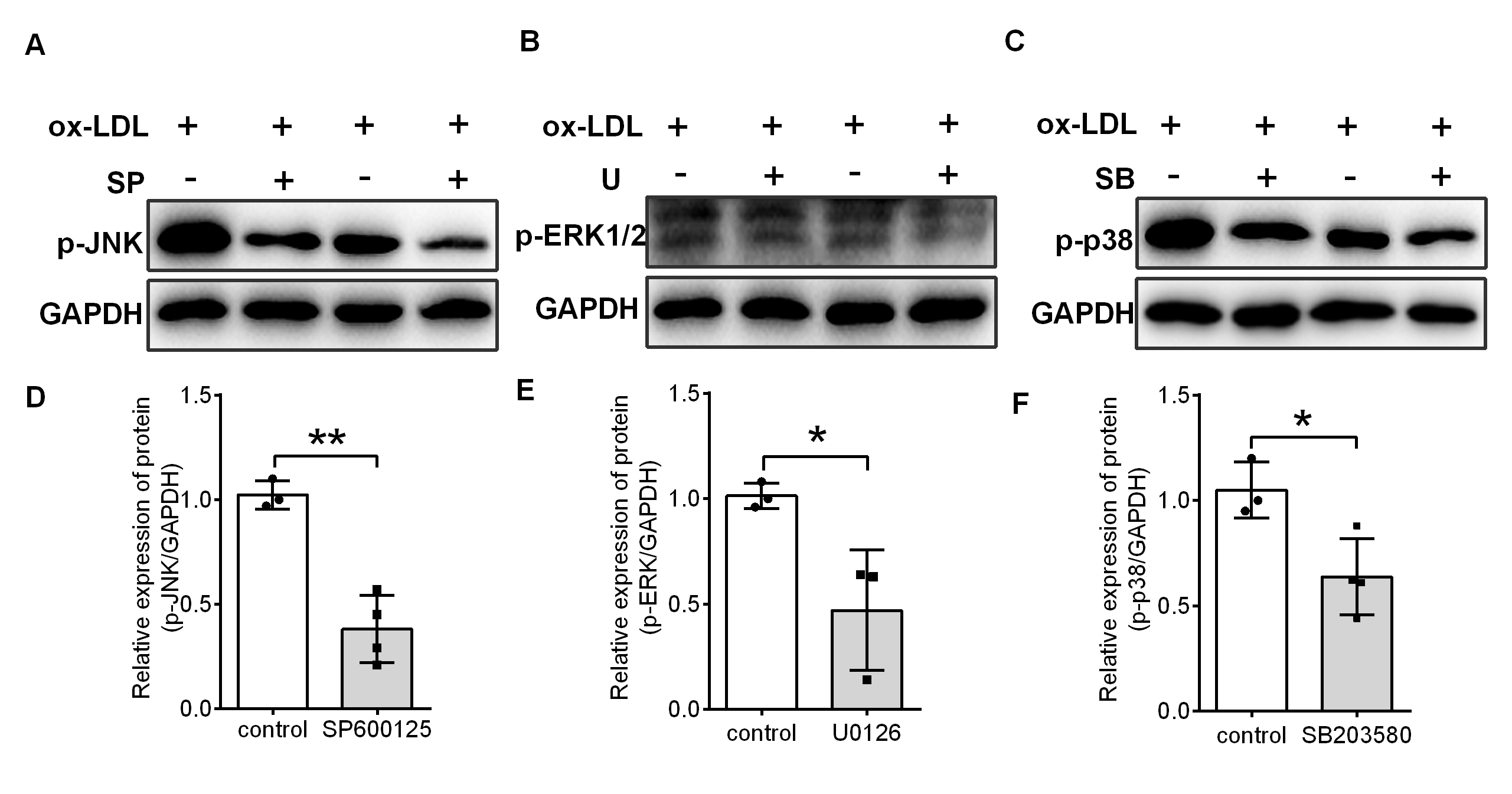

**Supplemental Figure 4**

(A) Representative images of the expression of p-JNK by western blot after the Raw264.7 macrophage cell line was treated with the p-JNK inhibitor SP600125 (10 μM) for 3 h, and ox-LDL (50 μg/ml) was added for 48 h. (B) Quantitative data from (A). (C) Representative images of the expression of p-p38 by western blot after the Raw264.7 macrophage cell line was treated with the p-p38 inhibitor SB203580 (10 μM) for 3 h, and ox-LDL (50 μg/ml) was added for 48 h. (D) Quantitative data from (C). (E) Representative images of the expression of p-ERK1/2 by western blot after the Raw264.7 macrophage cell line was treated with the p-ERK1/2 inhibitor U0126 (10 μM) for 3 h, and ox-LDL (50 μg/ml) was added for 48 h. (F) Quantitative data from (E). (G) Representative images of the expression of Irgm1 in macrophages as assessed by western blotting in si-control (Lane 1–5) and Si-Irgm1 (Lane 6–10) groups. (B, D, E, G) GAPDH was used as a loading control. Quantitative data represent the fold change after normalizing the band intensity of p-JNK, p-p38, p-ERK1/2, and Irgm1 to GAPDH. **p*<0.05, ***p*<0.01, ****p*<0.001. Results are presented as mean ± SD. Statistical analysis: unpaired Student’s *t-test*.

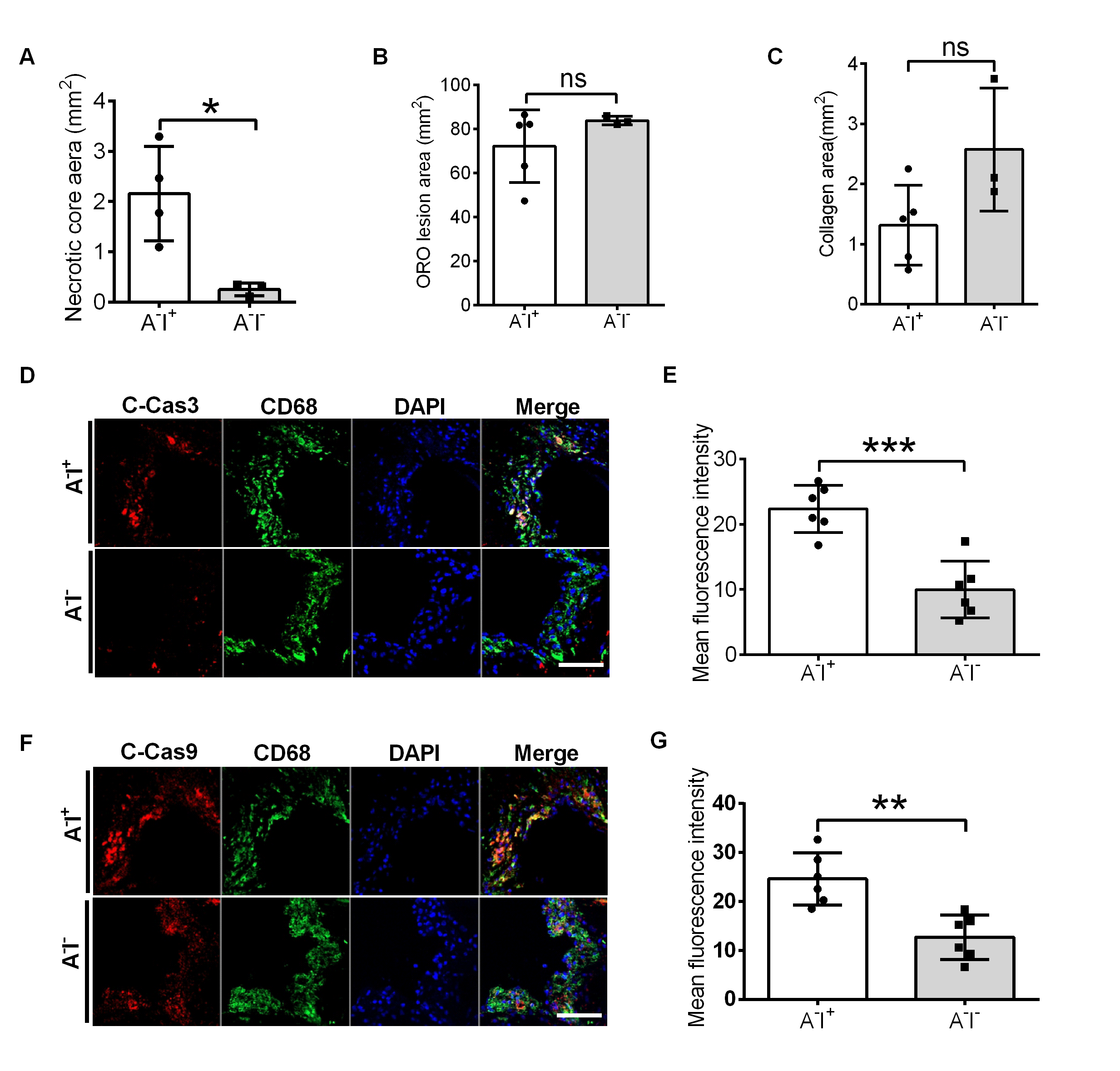

**Supplemental Figure 5**

(A–C) Quantitative data of the necrotic core area, ORO positive area, and collagen area in the aortic sinus plaque from ApoE^-/-^Irgm1^+/+^ (n=5) and ApoE^-/-^Irgm1^+/-^ mice (n=3). (D, E) Representative images (D) and quantitative data (E) for co-location of cleaved-caspase3 (red) and CD68 (green) by immunofluorescence staining after 16 weeks of a high-fat diet in ApoE^-/-^Irgm1^+/+^ (n=3) and ApoE^-/-^Irgm1^+/-^ bone marrow chimera recipient mice (n=3); nuclei were stained with DAPI (blue). (F, G) Representative images (F) and quantitative data (G) for co-location of cleaved-caspase9 (red) and CD68 (green) by immunofluorescence staining after 16 weeks of a high-fat diet in ApoE^-/-^Irgm1^+/+^ (n=3) and ApoE^-/-^Irgm1^+/-^ bone marrow chimera recipient mice (n=3); nuclei were stained with DAPI (blue). ApoE^-/-^Irgm1^+/+^ vs. ApoE^-/-^Irgm1^+/-^, **p*<0.05, ***p*<0.01, ****p*<0.001, ns=no significance. Scale bar=50 μm. Results are presented as mean ± SD. Statistical analysis: unpaired Student’s *t-test*.

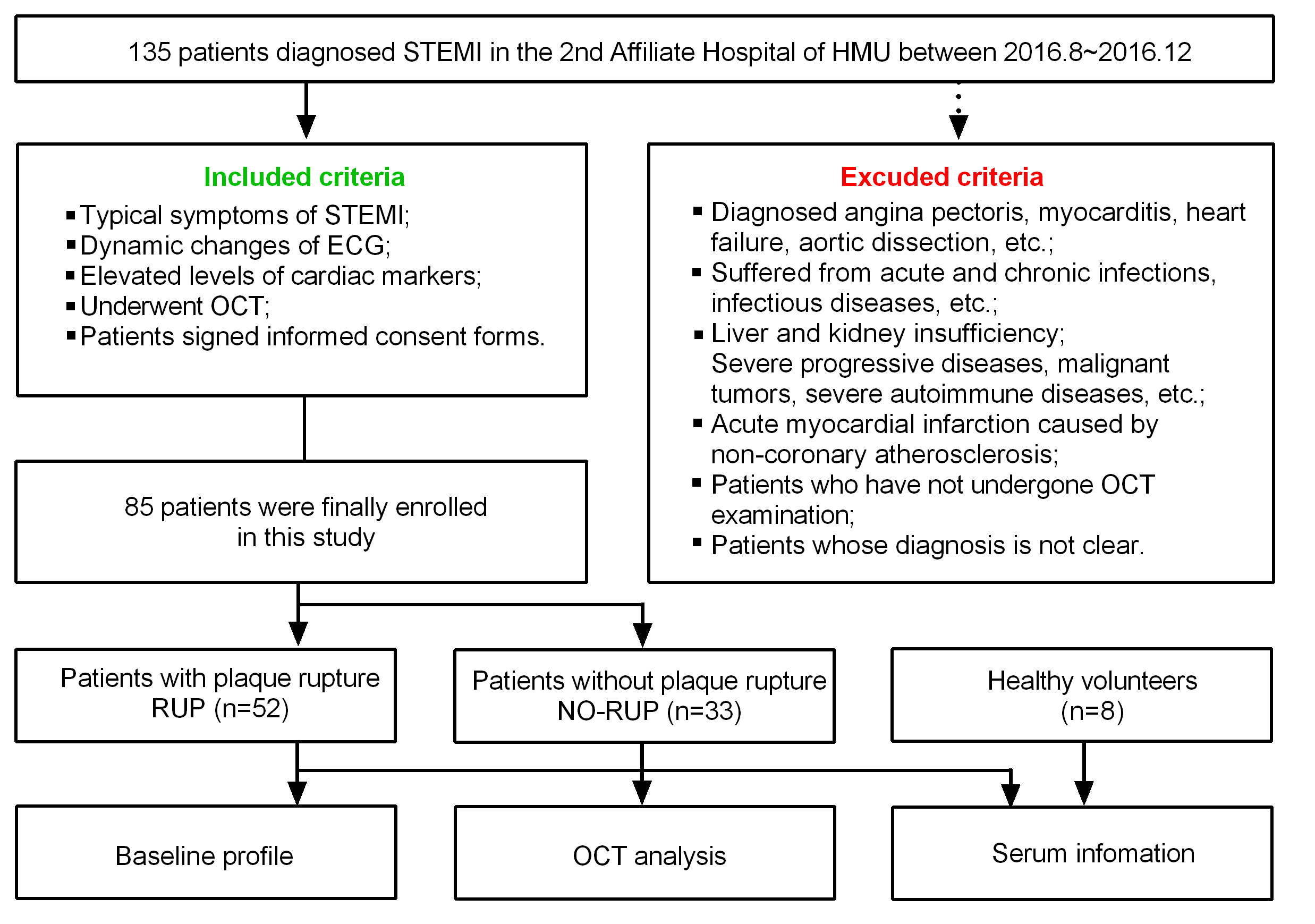

**Supplemental Figure 6. Study flow chart.** STEMI, ST segment elevation myocardial infarction; HMU, Harbin Medical University; ECG, electrocardiogram; OCT, optical coherence tomography.

**Supplemental Table 1. Clinical characteristics of patients for the present study**

| Covariate | RUP | NO-RUP | P value |
| --- | --- | --- | --- |
| No. of subjects | 52 | 33 |  |
| Age, yrs | 56.08±9.83 | 51.15±12.70 | 0.063 |
| Male | 39(75.0%) | 27(81.8%) | 0.462 |
| DM | 15(28.8%) | 6(18.2%) | 0.267 |
| Smoking | 30(57.7%) | 25(75.8%) | 0.089 |
| Hypertension | 30(57.7%) | 13(39.4%) | 0.301 |
| Hyperlipidemia | 22(42.3%) | 15(45.5%) | 0.776 |
| LDL-C, mg/dl | 3.29±0.90 | 2.82±0.94 | 0.039* |
| HDL-C, mg/dl | 1.25±0.36 | 1.21±0.37 | 0.635 |
| TG, mg/dl | 1.79±0.79 | 1.56±0.84 | 0.258 |
| TC, mg/dl | 4.92±0.87 | 4.37±0.98 | 0.019* |
| hs-CRP, mg/dl | 7.32±5.03 | 6.98±5.34 | 0.789 |
| Peak-TnI | 107.25±16.95 | 103.42±25.64 | 0.897 |

Values are expressed as Mean ± SD or n (%);

DM, diabetes mellitus; LDL-C, low-desity lipoprotein cholesterol; HDL-C, high-desity lipoprotein cholesterol; TG, triglycerides; TC, total cholesterol; hs-CRP, high sensitivity C-reactive protein; TnI, troponin I.

**Supplemental Table 2. OCT findings of the patients for the present study.**

| Covariate | RUP | NO-RUP | P value |
| --- | --- | --- | --- |
| No. of subjects | 52 | 33 |  |
| TCFA | 37(71.2%) | 4(12.1%) | <0.001* |
| Min FCT, mm | 0.05±0.02 | 0.07±0.03 | 0.026* |
| Lipid-rich plaque | 46(88.5%) | 12(36.4%) | <0.001* |
| Thrombus | 48(92.3%) | 31(93.9%) | 0.775 |
| Calcification | 8(15.4%) | 2(6.1%) | 0.194 |
| Cholesterol crystals | 18(34.6%) | 7(21.9%) | 0.215 |

Values are expressed as Mean ± SD or n (%);

TCFA, thin-cap fibroatheroma; FCT, fibrous cap thickness.

**Supplemental Table 3. Reagents or resources**

| **REAGENT OR RESOURCE** | **SOURCE** | **IDENTIFIER** |
| --- | --- | --- |
| NAC | Beyotime | Cat#:S0077 |
| ox-LDL | yiyuanBiotech | Cat#:YB-002 |
| SB203580 | Selleck | Cat#:S1076 |
| U0126 | Selleck | Cat#:S1102 |
| Lipofectamine3000 | Invitrogen | Cat#:L3000008 |
| Mounting-medium | Dako | Cat#:CS703 |
| SYBR Green qPCR Master Mix | MCE | Cat#:HY-K0501 |
| SDS-PAGE | Beyotime | Cat#:P0015 |
| PMSF | Beyotime | Cat#:ST505 |
| Phosphatase inhibitor | Bimake | Cat#:B15001-A |
| Hypersensitive ECL chemiluminescencekit | HaiGene | Cat#:M2301 |
| HE dye | Thermo | Cat#:67-63-0 (3%-5%) |
| Modified Oil Red O | Solarbio | Cat#:G1261 |
| Modified masson’s trichrome stain | Solarbio | Cat#:G1345 |
| DHE fluorescence probe | Beyotime | Cat#:S0063 |
| Skim milk powder | Biofroxx | Cat#:1172GR100 |
| Albumin Bovine V | Biotopped | Cat#:MW68000 |
| Triton X-100 | Biofroxx | Cat#:1139ML100 |
| DAPI | Beyotime | Cat#:C1002 |
| TRIzol | HaiGene | Cat#:B0201 |
| BCA | Beyotime | Cat#:P0012 |
| Dual color prestained protein marker | EpiZyme | Cat#:WJ101 |
| Caspase-3 colorimetric assay kit | Solarbio | Cat#:BC3830 |
| Caspase-9 colorimetric assay kit | Solarbio | Cat#:BC3890 |

**Supplemental Table 4. Antibodies**

| **ANTIBODIES** | **SOURCE** | **IDENTIFIER** |
| --- | --- | --- |
| LC3 | Sigma | Cat#:L7543 |
| F4/80 | Biolegend | Cat#:123113 |
| CD11b | BD | Cat#:553312 |
| p62 | MBL | Cat#:PM045 |
| cleaved-caspase-3 | CST | Cat#:ASP175 |
| cleaved-caspase-9 | CST | Cat#:ASP353 |
| GAPDH | ZSGB-BIO | Cat#:ta-08 |
| p-ERK1/2 | Wanleibio | Cat#:WL02368 |
| CD68 | Abcam | Cat#:ab53444 |
| CD68 | ZSGB-BIO | Cat#:ZM-0464 |
| p-JNK | CST | Cat#:9251S |
| p-p38 | CST | Cat#:4092S |
| bax | Abcam | Cat#:ab182733 |
| bcl2 | Abcam | Cat#:ab18858 |
| IRGM | AbMart | Cat#:Clone:IG9 |
| Irgm1 | AbMart | Cat#:Clone:IC11 |
| TRITC Goat Anti-Rabbit IgG H&L | Abcam | Cat#:ab6719 |
| FITC goat anti-mice IgG | Abcam | Cat#:ab6785 |
| FITC goat anti-rat | ZSGB-BIO | Cat#:ZF0315 |
| FITC Annexin V | BD | Cat#:556547 |

**Supplemental Table 5.**

| **Gene** | **Forward primers (5’-3’)** | **Reverse primers (5’-3’)** |
| --- | --- | --- |
| Atg5 | CTTCTGCACTGTCCATCTAAGG | ATCCAGAGTTGCTTGTGATCTT |
| Atg7 | GCAGCCAGCAAGCGAAAG | CCGGTCTCTGGTTGAATCTCCTG |
| Beclin 1 | CCCGTGGAATGGAATGAGATTA | CCGTAAGGAACAAGTCGGTATC |
| LC3 | AGAGTGGAAGATGTCCGGCT | CACTTCGGAGATGGGAGTGG |

**Supplemental Table 6.**

| Gene | primers (5’-3’) |
| --- | --- |
| LRG1 | GGAGAAAGTGAAGTACCC |
| LRG2 | CTCTGACACCGAGAGAAT |
| Neo3 | CATTTGTCACGTCCTGCA |
